# The MutRS quorum sensing system controls lantibiotic mutacin production in the human pathogen *Streptococcus mutans*

**DOI:** 10.1101/2024.10.01.616031

**Authors:** Ryan M. Wyllie, Paul A. Jensen

## Abstract

Microbes use quorum sensing systems to respond to ecological and environmental changes. In the oral microbiome, the pathogenic bacterium *Streptococcus mutans* uses quorum sensing to control the production of bacteriocins. These antimicrobial peptides kill off ecological competetors and allow *S. mutans* to dominate the microenviornment of dental plaques and form dental caries. One class of bacteriocins produced by *S. mutans*, the lantibiotic mutacins, are particularly effective at killing due to their broad spectrum of activity. Despite years of study, the regulatory mechanisms governing production of lantibiotic mutacins I, II, and III in *S. mutans* have never been elucidated. We discovered a distinct class of quorum sensing system, MutRS, that regulates mutacins and is widespread among the streptococci. We demonstrate that MutRS systems are activated by a short peptide pheromone (Mutacin Stimulating Peptide, or MSP) and show that MutRS controls production of three separate lantibiotic mutacins in three different strains of *S. mutans*. Finally, we show that paralogous MutRS systems participate in inter- and intrastrain crosstalk, providing further evidence of the interplay between quorum sensing systems in the oral streptococci.

## 1 Introduction

Dental caries, also known as tooth decay or dental cavities, is the most common chronic disease in humans, affecting over two billion adults and 500 million children worldwide [1]. Members of the oral streptococci, specifically *Streptococcus mutans*, are the primary etiological causes of dental caries [2]. However, in recent years our view of cariogenesis has shifted from a focus on individual species towards the dynamics of microbial communities [3], [4]. The oral microbiome is highly diverse in healthy individuals and composed of more than 600 bacterial and 100 fungal species [5]. Occasionally, a small number of acid-tolerant and acid-producing streptococci come to dominate the tooth microenviornment. These bacteria produce excessive amounts of lactic acid as they grow, which demineralizes the tooth enamel and ultimately leads to the formation of dental caries [6].

The community-level interactions that lead to oral dysbiosis remain poorly understood despite their importance to human health. Our lack of understanding is largely due to the high dimensionality of the interaction space. Interactions between oral microbes can arise from adhesion requirements [7]–[9], metabolic cross-talk [10], [11], pH modulation [12]–[14], oxidative stressors [15], [16], and cooperative biofilm development [17]. The oral streptococci also interact by producing bacteriocins— ribosomally synthesized and post-translationally modified peptides (RiPPs) with antimicrobial activity that are weapons for both the pathogenic and the commensal streptococci [18]. Bacteriocins frequently target closely related species that occupy and compete within the same ecological niche as the producer strain. However, the lantibiotics subclass of bacteriocins, also known as Class I bacteriocins, have a broad spectrum of activity and can induce large perturbations to the oral microbiome [19].

Lantibiotic bacteriocins contain cyclical domains bridged by the non-proteinogenic amino acids mesolan-thionine and 3-methyllanthionine [18], [20]. These amino acids are the result of extensive modification by enzymes typically encoded within the same operon as the lantibiotic precursor gene and are crucial for antimicrobial activity and molecular stability [19]. The bacteriocins of *S. mutans*, known as mutacins, include several lantibiotics: mutacins I, II, III, 1140, K8, and Smb. Given their broad spectrum of activity [18], lantibiotic mutacin production confers a major competitive advantage to *S. mutans* within the plaque microenviornment [21]. Unsurprisingly, lantibiotic operons have been found in the genomes of several virulent clinical isolates from carious lesions [22]–[24].

The regulation of lantibiotic mutacin production in *S. mutans* has remained poorly understood since their initial characterization over three decades ago [22]–[24]. The non-lantibiotic mutacins and the two-peptide lantibiotic mutacin Smb are known to be directly regulated by the ComCDE quorum sensing system [19], [25], [26], with secondary control exerted by the HdrR and BrsR systems [27], [28] and indirect regulation through the quorum sensing system ComRS [29], [30]. However, these systems do not appear to directly regulate expression of the majority of lantibiotic mutacin genes. Shortly after the initial discovery of the lantibiotic mutacins, a gene named *mutR* encoding an “Rgg” homolog was identified upstream of the mutacin I, II, III, and 1140 operons [22], [23], [31]. In subsequent studies, *mutR* was shown to be necessary for lantibiotic mutacin gene expression [31]. However, relatively constant levels of *mutR* transcript were observed despite large variations in lantibiotic gene expression, suggesting that lantibiotic gene expression was not controlled at the level of *mutR* transcription [32]. No other regulators of the mutacin I, II, III, and 1140 operons have been identified.

Many bacteriocin genes (including the non-lantibiotic mutacins) are regulated by quorum sensing (QS) systems [33]–[36]. The QS systems of the oral streptococci and many other Gram-positive bacteria use short, genomically-encoded oligopeptide pheromones as autocrine signalling molecules. The pheromones are typically translated in an inactive form with a variable-length leader sequence targeting the nascent peptide to processing and export machinery. Once in the extracellular space, the pheromone accumulates proportionally to the abundance of pheromone-producing cells until a critical concentration (“quorum”) is met. The pheromones signal back into the responsive cells and activate transcriptional rewiring through a variety of mechanisms [29], [37]–[39].

Here, we show that MutR is not, as previously thought, an Rgg-class regulator. Instead, the MutR regulators form a distinct class of QS systems. We identified the cognate peptide MutS for several MutR regulators. The mature form of the MutS pheromone, which we name Mutacin Stimulating Peptide (MSP), regulates mutacin I, II, and III production in three strains of *S. mutans*. Furthermore, we show that closely related but distinct MutRS systems are capable of crosstalk at the peptide-binding and DNA-binding levels.

## 2 Results

### 2.1 MutRS represents a distinct class of peptide-based QS systems

QS systems regulate several streptococcal bacteriocin gene clusters, so we hypothesized that MutR was the regulator protein of an uncharacterized QS system. To identify a putative pheromone gene, we compiled a database of characterized streptococcal Rgg-Shp and ComRS QS systems as well as MutR regulators from *S. mutans*. We then expanded our database by homology searches using known regulators from other streptococci. We manually predicted the peptide pheromone gene and regulator DNA binding motif for each regulator. Our final database includes 483 streptococcal QS systems from 179 strains of 59 streptococcal species (Supplementary Data 1). The database includes predicted or previously validated pheromones and DNA binding motifs for 327 (67.7%) of the QS systems.

We used the NCBI COBALT tool to visualize the phylogenic relationships between the regulators (Figure 1A). Three major regulator clusters emerged. The largest cluster includes 260 Rgg-Shp systems. Generally, *shp* pheromone genes are found upstream of and divergent from their regulator gene. Mature SHP pheromones typically contain an N-terminal acidic residue followed by a stretch of 6–12 hydrophobic and largely aliphatic resides. Binding motif sequences varied considerably across the entire Rgg cluster but were conserved within smaller sub-clusters. However, we typically found Rgg binding motifs ∼25 nucleotides upstream of a −10 element, followed by variable spacing before the *shp* Shine-Delgarno Sequence (SDS) (Figure 1B).

**Figure 1:**
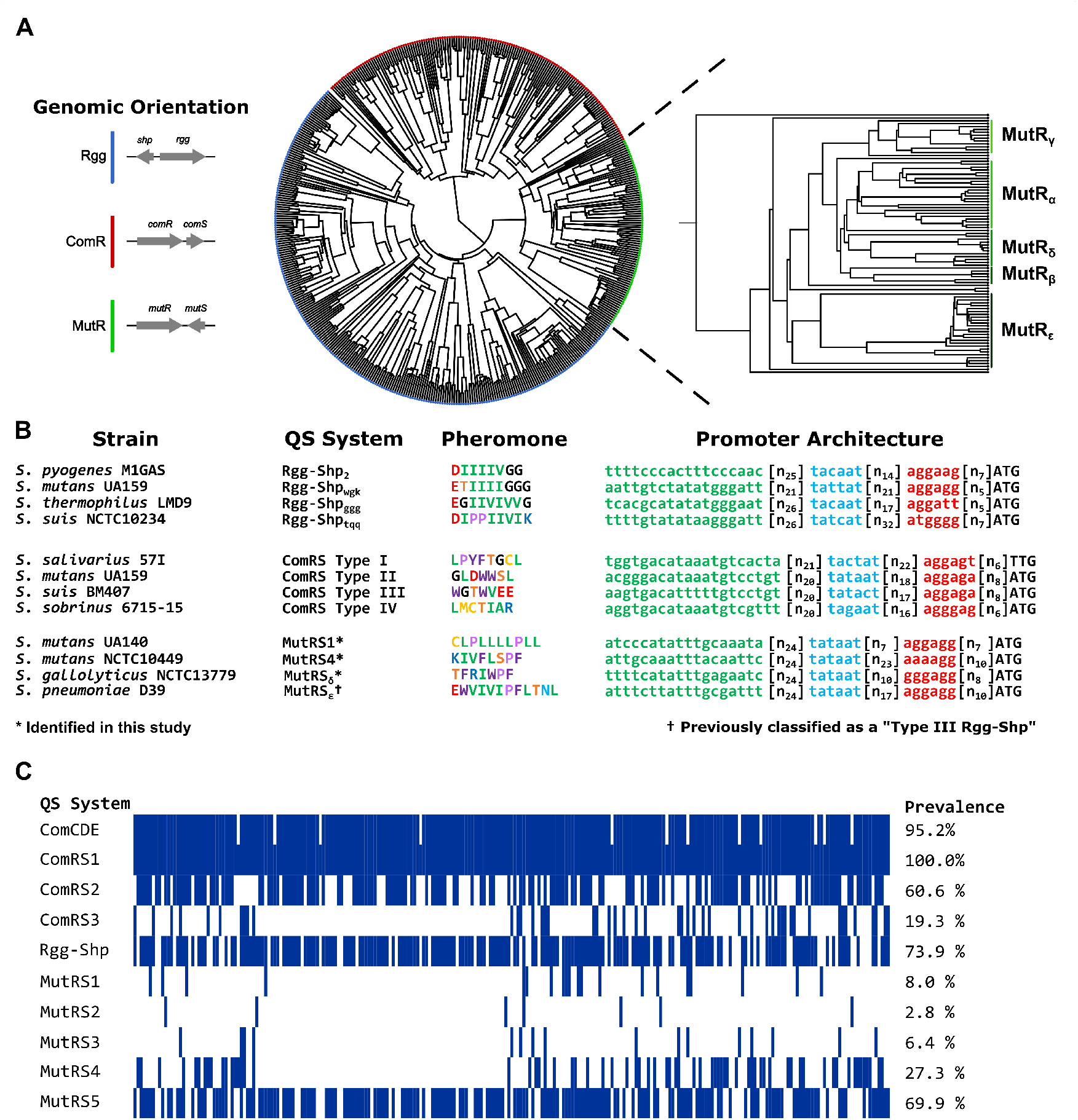
MutRS forms a separate class of streptococcal quorum sensing systems. **A**. MutR transcriptional regulators are homologous to, but distinct from, other peptide-based quorum sensing systems like the Rgg and ComR families. **B**. The Rgg, ComR, and MutR regulator families are distinguished by 1. the orientation of the regulator and pheromone genes, 2. the primary sequence of the mature pheromones, and 3. the promoter architecture of each system’s cognate peptide gene. Four representative pheromones from each class are colored according to their biochemical properties using the RasMol color scheme [40]. The promoters for these pheromones are also shown with the putative regulator binding site in green, the −10 promoter element in blue, and the pheromone gene’s Shine-Delgarno Sequence (SDS) in red. The start codon for each pheromone gene is capitalized. **C**. A collection of 249 *S. mutans* genomes contain different peptide-based QS systems. MutRS1, MutRS2, and MutRS3 systems represent the MutRS_*α*_ subcluster, while MutRS4 and MutRS5 come from the MutR_*β*_ and MutR_*γ*_ subclusters, respectively.

The second largest cluster contains 137 ComRS systems. For these systems the *comS* pheromone gene is found downstream of and on the same strand as the regulator gene. ComRS pheromones vary widely in their biochemical properties, unlike the conserved Rgg-Shp pheromone properties described above. However, the promoters of dissimilar *comS* genes are similar. We typically found a ComR binding site (5’-GACANNNNTGTC-3’) ∼24 nucleotides upstream of a −10 element and the *comS* SDS (Figure 1B).

To our surprise, the third major cluster (*n* = 86) was composed entirely of MutR-type regulators. Importantly, the predicted cognate pheromone genes of MutRS systems, which we call *mutS*, are found downstream of and in a convergent orientation with their regulator genes (Figure 1A). As with the ComRS cluster, the predicted MutS-derived pheromone sequences have varied biochemical properties. In general, the predicted mature MutS-derived pheromones have a N-terminal hydrophilic residue followed by a hydrophobic stretch with frequent prolines and bulky aromatic resides. MutR binding motif sequences varied across the entire MutRS cluster but were conserved within five subclusters. We refer to these MutRS subclusters by Greek letter suffixes (*α*–*ε*). For MutRS systems, the *mutS* promoter contains a predicted MutR binding motif ∼ 24 nucleotides upstream from a −10 element that is in turn variably spaced to the SDS of *mutS* (Figure 1B). We call this MutR binding motif a “MutR box”. Overall, MutRS systems are distinct from Rgg-Shp and ComRS QS systems by protein homology, pheromone-regulator genomic orientation, pheromone primary sequence, and regulator DNA binding motifs.

Next, we analyzed the pangenome of *S. mutans* to estimate the prevalence of MutRS and other pheromone-based QS systems. We searched 249 publicly available *S. mutans* genomes for distinct regulators (<85% intraclass pairwise sequence identity to regulators from our database, Supplementary Data 2). Most of the regulators in our database are cytoplasmic and directly bind to cognate pheromones, but we also included regulators from the two-component ComCDE systems (Figure 1C). Unsurprisingly, the two best characterized *S. mutans* QS systems, ComCDE and ComRS1, are present in 95.2% and 100% of sequenced *S. mutans* strains, respectively. Two recently characterized QS systems, an Rgg-Shp system [39] and the ComRS2 system [41], are also widespread. Interestingly, we identified a third, unstudied ComRS-type system in nearly 20% of *S. mutans* strains. Three of the five MutRS subclusters identified in Figure 1A are found in the *S. mutans* pangenome: MutR_*α*_, MutR_*β*_, and MutR_*γ*_. We predicted cognate phermomone genes and regulator binding sites for all three clusters. Overall, 88.0% of *S. mutans* strains contain at least one MutR-class regulator, while 23.7% contain two or more MutR homologs. Thus, MutR regulators appear almost as frequently as ComRS or Rgg-Shp systems in *S. mutans*.

With our bioinformatic survey complete, we set out to validate our predictions regarding the MutR_*α*_ regulators associated with *S. mutans* lantibiotic mutacin operons. The twenty seven MutRa regulators in our QS database were found in a diverse group of streptococci including *S. mutans, S. sobrinus, S. gordonii*, and *S. thermophilus* (Supplementary Figure S1). Interestingly, MutR_*α*_ regulators are found upstream of both lantibiotic and non-lantibiotic bacteriocin operons. At least one of three distinct MutR_*α*_ regulators can be found in ∼17% of all *S. mutans* strains (Figure 1C). We refer to these *S. mutans* MutR regulators as MutR1, MutR2, and MutR3. These three regulators have ∼60% pairwise sequence identity and were found within distinct gene clusters. MutR1 regulators appear upstream of the lantibiotic mutacin I, III, and 1140 operons, while MutR2 regulators appear upstream of the lantibiotic mutacin II operon (Figure 2A). MutR3 is flanked on both sides by several transposon-associated genes, making it unclear what (if any) genes it regulates. Because of ambiguity in the MutR3 regulon, we focused on characterizing the MutR1 and MutR2 regulators controlling the mutacin I, II, and III systems (Figure 2A). We note that the MutR1 regulator and the MutS1 peptide encoded upstream of the mutacin 1140 operon in *S. mutans* strain ATCC55676 are identical to those associated with the mutacin III operon in *S. mutans* strain LAR01.

**Figure 2:**
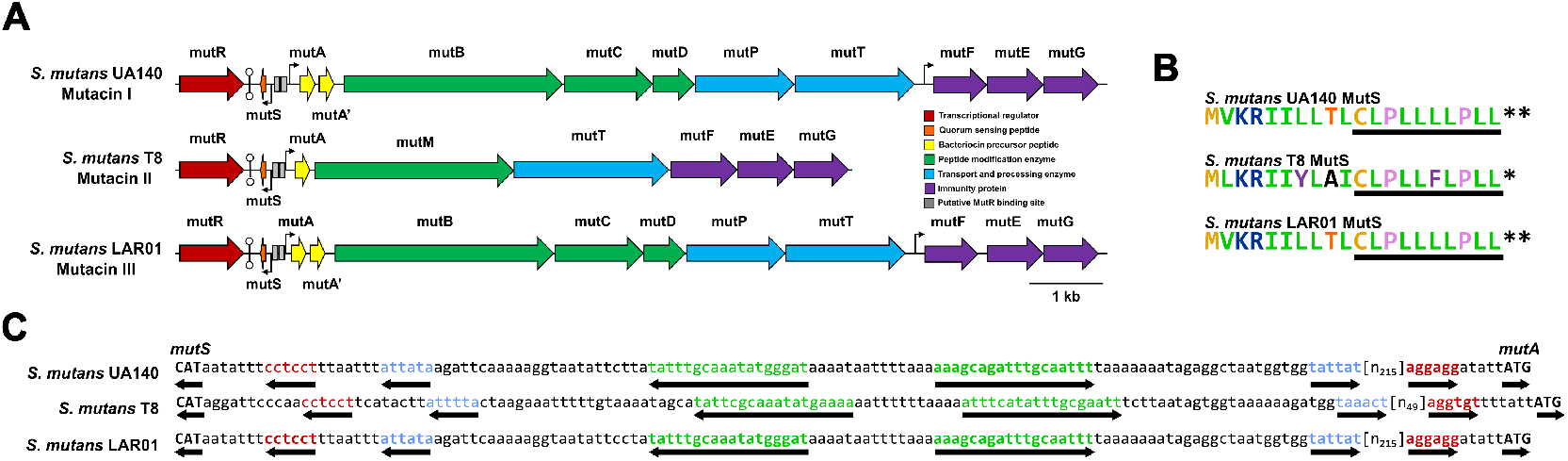
MutRS systems regulate three different lantibiotic mutacin operons in *S. mutans*. **A**. The mutacin I operon in *S. mutans* strain UA140, the mutacin II operon in *S. mutans* strain T8, and the mutacin III operon in *S. mutans* strain LAR01 have a MutRS system upstream of lantibiotic biosynthesis, transport, and immunity genes. A duet promoter upstream of the lantibiotic precursor *mutA* and the quorum sensing pheromone gene *mutS* is responsible for transcription of these genes, while in strains UA140 and LAR01 immunity genes are also transcribed from a second promoter upstream of *mutF*. Light grey boxes indicate the predicted mutR binding sites. In all strains, a bidirectional rho-independent transcriptional terminator was predicted to insulate the *mutR* gene from the *mutS* gene. **B**. The primary sequences of the MutS peptides show a basic-hydrophobic leader sequence followed by a helix-breaking cysteine residue. The predicted mature pheromone sequence is underlined. **C**. The duet promoter architecture governing transcription of *mutS* and the lantibiotic operon is shown for the mutacin I, II, and III operons. Start codons for *mutS* and *mutA* are capitalized and bolded. The predicted SDS, −10 promoter elements, and MutR binding motifs are shown in red, blue and green, respectively.

### 2.2 MutS-derived peptides activate lantibiotic mutacin gene expression in *S. mutans*

The MutR1 regulator encoded upstream of the mutacin I operon in *S. mutans* UA140 has 97.9% protein sequence similarity to the MutR1 regulator of mutacin III in *S. mutans* LAR01. The two regulators differ only at residue 83 (asparagine in UA140 and histidine in LAR01) and a five amino acid truncation at the C-terminal end of the LAR01 regulator. By contrast, the MutR2 regulator encoded upstream of the mutacin II operon in *S. mutans* T8 shares 61% amino acid identity and 78% sequence similarity to the MutR1s from UA140 and LAR01. Mirroring this difference in protein homology, the MutS2 peptide differs from the MutS1 peptide. The MutS2 peptide features two substitutions in the leader sequence and a leucine to phenylalanine substitution in the putative mature pheromone sequence (Figure 2B). Despite peptide sequence differences, the promoter regions for these *mutS* genes all feature a predicted duet architecture with one MutR-controlled promoter driving expression of the *mutS* gene, and another MutR-controlled promoter driving expression of the lantibiotic precursor gene *mutA* and the rest of the lantibiotic operon (Figure 2C).

We built a dual fluorescent reporter cassette to determine if the predicted mature pheromone peptides activate lantibiotic mutacin production in *S. mutans*. We inserted the reporter cassette into a transcriptionally insulated locus on the chromosomes of each *S. mutans* strain. The DNA sequence between the start codons of each strain’s *mutS* and *mutA* genes was inserted between two spectrally distinct and rapidly-maturing fluorescent proteins: mScarlet and mNeonGreen (Figure 3A). Mid log-phase cultures of the reporter strains were then dosed with either a DMSO control or a synthesized peptide consisting of the last nine, ten, or eleven C-terminal amino acids of the respective MutS peptide. DMSO controls failed to produce fluorescence above background, while MutS-derived peptides resulted in activation of P_*mutS*_ and P_*mutA*_ in all three strains (Figure 3B).

**Figure 3:**
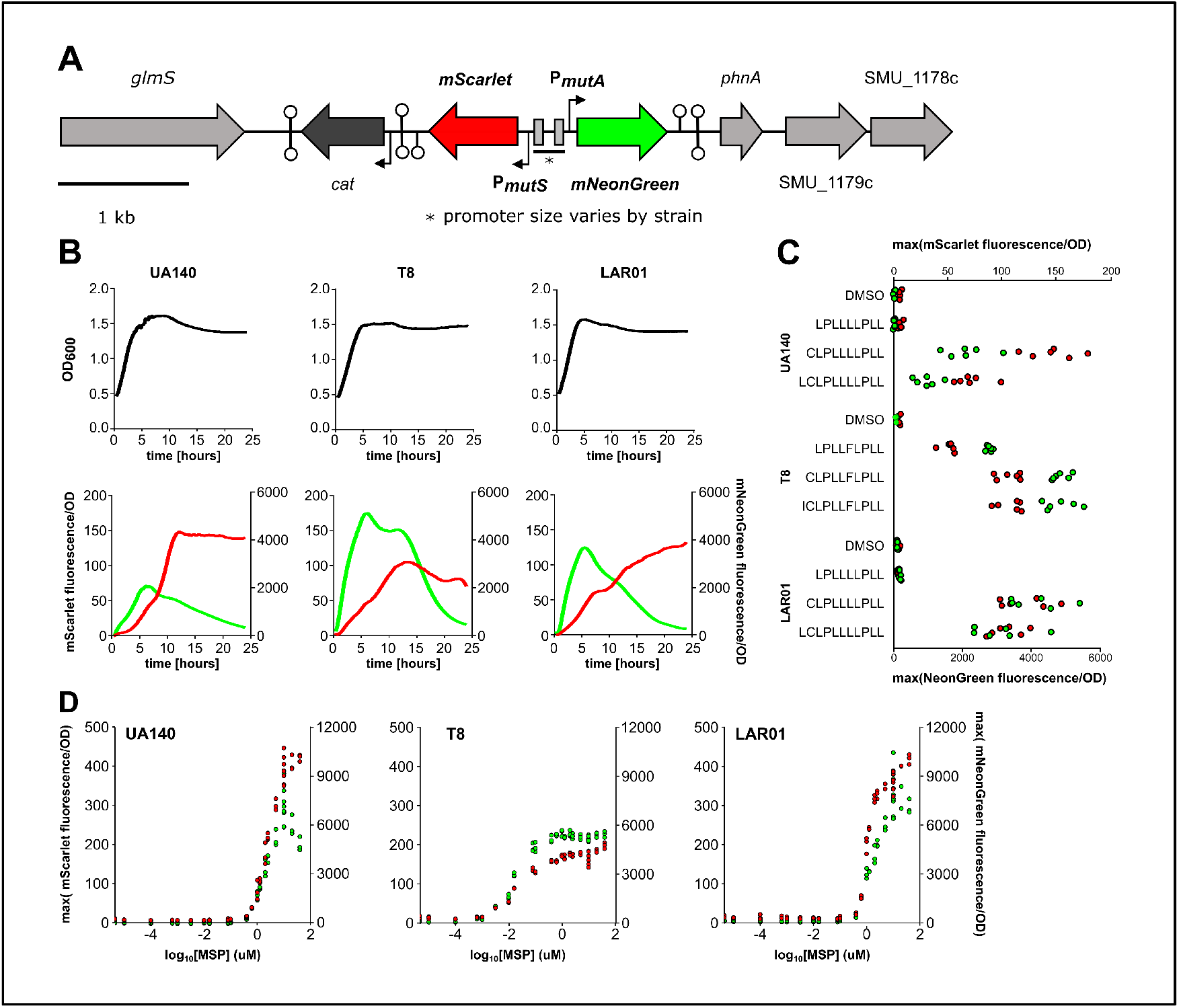
Lantibiotic gene activation levels are a function of pheromone concentration.**A**. We built a dual fluorescent reporter cassette and integrated it into a defined locus on the *S. mutans* chromosome. P_*mutS*_ activity and P_*mutA*_ activity are detected by measuring the abundance of the fluorescent proteins mScarlet-I and mNeonGreen, respectively.**B**. A representative time series readout for lantibiotic reporter strains dosed with their respective ten amino acid long, MutS-derived pheromone. Addition of the pheromone immediately coactivates P_*mutS*_ and P_*mutA*_ in all three strains, consistent with the predicted duet promoter architecture.**C**. The C-terminal ten amino acids make up the mature Mutacin Stimulating Peptide (MSP). Truncated nine amino acid pheromone variants failed to activate transcription at P_*mutS*_ and P_*mutA*_ in *S. mutans* UA140 and LAR01; while resulting in greatly attenuated activation in *S. mutans* T8.**D**. Transcriptional activation of P_*mutS*_ and P_*mutA*_ occurs in an MSP dose-dependent manner. The MutR1 systems found in UA140 and LAR01 have a higher maximal activation but a lower sensitivity to MSP1. The MutR2 system from T8 is highly sensitive to MSP2 levels but results in a lower maximal activation relative to the MutR1 sytems.

For all three *S. mutans* strains, the ten or eleven amino acid peptide activated both P_*mutS*_ and P_*mutA*_ (Figure 3C). Activity of P_*mutA*_ was 75-fold higher than P_*mutS*_ in strain T8 and 20-fold higher in UA140 and LAR01, indicating that the lantibiotic operon promoter is stronger than the pheromone gene promoter. In UA140 and LAR01, the nine amino acid peptide failed to generate signal above background, while in T8 the nine amino acid peptide resulted in half the activation achieved by the longer peptides. The N-terminal cysteine at residue ten is thus important for binding and/or activation of MutR1 and MutR2. We used the ten amino acid peptide for all subsequent experiments and refer to it as Mutacin Stimulating Peptide (MSP); however, we do not know the exact location of the MutS cleavage site that produces active MSP *in vivo*. We refer to the UA140 and LAR01 MSP as MSP1 and the T8 MSP as MSP2.

In *S. mutans* T8, we predict that the immunity genes for the mutacin II operon are cotranscribed with the precursor peptide *mutS* and biosynthetic enzyme genes. However, an intergenic region of ∼200 bp separates the end of *mutT* and the start of*mutF* in the mutacin I and III operons (Figure 2A). To validate MSP activation of immunity gene expression, we constructed *mutF* reporter strains for *S. mutans* UA140 and LAR01. Adding MSP1 caused *mutR1*-dependent activation of P_*mutF*_ in both strains (Supplementary Figure S2). However, we failed to identify a nucleotide sequence in these strains that resembles the predicted MutR box found in the P_*mutS*_ and P_*mutA*_ regions. The lack of a MutR box suggests that MutR may bind other motifs or that immunity gene regulation may be controlled indirectly by MutR. Furthermore, no intrinsic terminator sequence was identified within the *mutT*–*mutF* intergenic region, so polycistronic transcription from the *mutA* promoter may supplement the transcriptional activity of the independent *mutF* promoter.

### 2.3 MSP activation of lantibiotic mutacin genes is a dose and *mutR*-dependent process

The *mutS* and *mutA* reporter activities increase with MSP concentration (Figure 3D). The minimum MSP concentrations that activated P_*mutS*_ and P_*mutA*_ were 0.4 *μ*M for UA140, 3.2 nM for T8, and 0.4 *μ*M for LAR01. The P_*mutS*_ signal increased through the highest concentration tested in all three strains, while the P_*mutA*_ signal reached a maximum value or plateaued at MSP concentrations of 1 *μ*M for T8 and 10 *μ*M for UA140 and LAR01.

Deleting *mutR* abolished the reporter signal in all surveyed conditions, indicating that MutR is necessary for MSP activation of both P_*mutS*_ and P_*mutA*_ (Figure 4). Supplementing the Δ*mutS* strains with 2 *μ*M exogenous MSP partially restored the wild-type promoter responses, reaching 73% and 53% of maximum activation for P_*mutA*_ and P_*mutS*_ in UA140, 56% and 30% of maximum activation in LAR01, and 94% and 60% in T8. Especially in T8, high levels of exogenous MSP can compensate for the loss of *mutS*, in part due to the higher sensitivity of T8’s MutRS2 system to MSP2 levels (Figure 3D).

**Figure 4:**
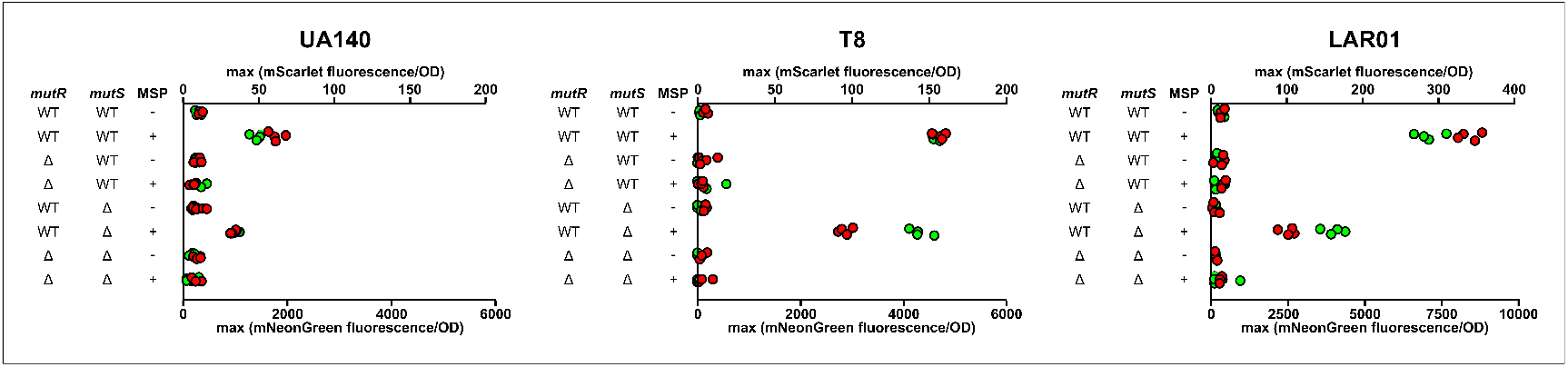
MSP activation of lantibiotic gene expression depends on *mutR* in three strains of *S. mutans*. Strains without *mutR* failed to activate transcription of lantibiotic genes, while *mutS* knockouts showed reduced activation. (WT = wild type. Δ = gene deletion strain. − = DMSO only. + = 2 *μ*M MSP.)

### 2.4 MSP activation of MutRS inhibits lantibiotic-sensitive strains

We used deferred antagonism assays to determine if MSP’s activation of lantibiotic genes resulted in the production of mature lantibiotic mutacins. Briefly, wild-type and Δ*mutRS* strains for UA140, T8, and LAR01 were spotted onto THY agar plates that had been supplemented with the appropriate MSP or a DMSO control. After allowing sufficient time for lantibiotic production, the plates were sterilized with UV irradiaton and an agar overlay containing one of five lantibiotic-sensitive strains [18], [42]–[45] was poured over the plates. Following overnight incubation, MSP-enhanced and *mutRS*-dependent zones of inhibition were observed for all three producer strains against at least one indicator strain (Figure 5). However, the inhibition profiles differed for mutacin I, mutacin II, and mutacin III-producing strains. *S. mutans* LAR01 and UA140 showed the broadest and narrowest spectrum of activity, respectively. Non-MutRS-dependent inhibition was observed for a number of producer-indicator pairings, suggesting that the producer strains contain other inhibitory mechanisms.

**Figure 5:**
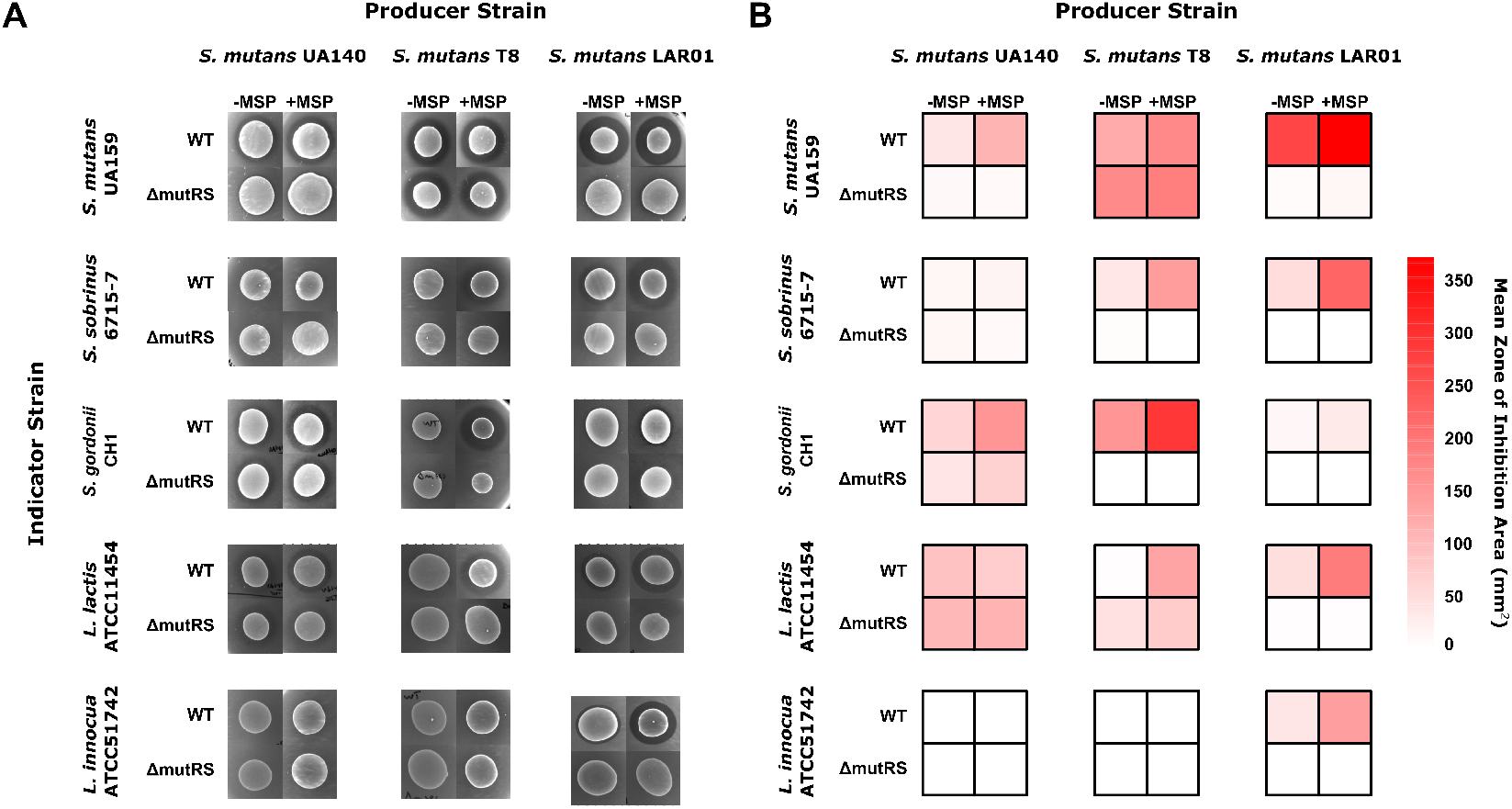
MSP’s activation of lantibiotic genes inhibits lantibiotic-sensitive strains, including species from the human oral cavity.**A**. Representative spots from a deferred antagonism assay show the effects of adding MSP and deleting MutRS. Wild type or *mutRS* knockout strains of *S. mutans* UA140, T8, and LAR01 were spotted onto THY agar supplemented with MSP or a DMSO control. Producer strains were grown overnight and killed by UV irradiation. Five lantibiotic-sensitive indicator strains were added as overlays: the *S. mutans* type strain UA159, an oral commensal *S. gordonii* CH1, a dental pathogen *S. sobrinus* 6715-7, a dairy lactic acid bacterium *Lactococcus lactis* ATCC11454, and a food-contaminating bacterium *Listeria innocua* ATCC51742. **B**. A heat map shows the mean area (*n*=3–4) for zones of inhibition in the deferred antagonism assays. The inhibitory profile of the three producing strains varies, but MSP-enhanced and *mutRS*-dependent inhibition was observed against at least one indicator species for all three producing strains.

### 2.5 MutRS systems participate in interstrain and intrastrain crosstalk

Crosstalk between closely related QS systems has been observed in Rgg-Shp and ComRS systems at both the peptide-binding [46], [47] and DNA-binding levels [41], [46]. We wanted to know whether similar crosstalk occurs between the three MutRS_*α*_ systems in *S. mutans*. However, care must be taken with studying MutRS3. The *mutR3* gene is flanked by repeat regions carrying transposon-associated genes and is an orphan contig in every published *S. mutans* genome except for LAR01. Interestingly, the *mutS3* gene, complete with the expected promoter architecture, is located 4 kilobases upstream of, and in a parallel orientation with the *mutR3* gene (Figure 6A). This is an exception to the genomic orientations of different QS systems shown in Figure 1A, although it is not unexpected given the transposon-mediated rearrangement. The colocalization of MutRS3 to a region with mobile genetic elements may impact MutRS3’s function in LAR01.

**Figure 6:**
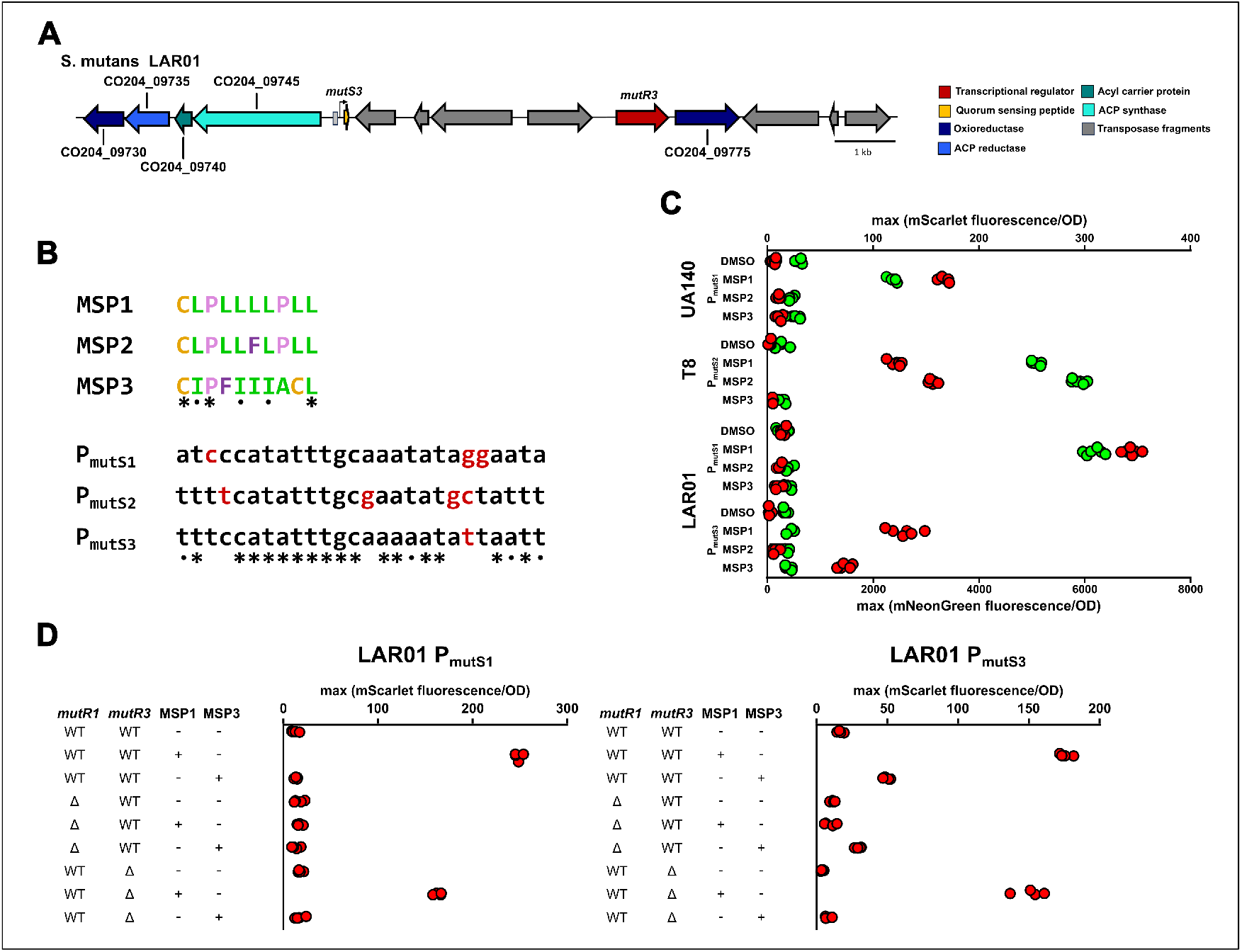
MutRS systems participate in inter- and intra-strain crosstalk. **A**. The *mutS3* pheromone and *mutR3* regulator genes in *S. mutans* LAR01 are separated by mobile genetic elements. We predict that MutRS3 regulates expression of a neighboring putative fatty acid biosynthesis operon. **B**. The primary amino acid sequence of MSP and the predicted P_*mutS*_ MutR-binding site suggest peptide-mediated and DNA binding-mediated MutRS interactions. An asterisk (*) indicates conserved amino acids or nucleotides across MutRS1, MutRS2, and MutRS3, while a period (.) indicates similar biochemical properties (hydrophobicity/sterics for MSP peptides or hydrogen bonding capacity for MutR DNA binding motifs.) **C**. Crosstalk occurs in a subset of possible MutR-MSP pairings. P_*mutS*_ reporter strains were dosed with MSP1, MSP2, and MSP3 to measure interactions between closely related MutRS_*α*_ systems. The MutRS1 systems in UA140 and LAR01 only respond to their own pheromones. The lone MutRS systems in T8 (MutRS2) responds to both MSP1 and MSP2, even though MSP1 is not found in the T8 genome. Strain LAR01 contains MutRS1 and MutRS3 systems, and the latter responds to both MSP1 and MSP3. **D**. The activity of P_*mutS1*_ and P_*mutS3*_ in Δ*mutR1* and Δ*mutR3* strains suggest that the crosstalk between the MutRS1 and MutRS3 systems is peptide-specific but DNA-binding promiscuous.

To identify crosstalk between the three MutRS systems, P_*mutS1*_,P_*mutS2*_, and P_*mutS3*_ reporter strains were grown to mid-log phase and dosed with MSP1, MSP2, or MSP3 (Figure 6C). The MutRS1 system in *S. mutans* UA140 only responds to its cognate pheromone, MSP1, but the MutRS2 system in *S. mutans* T8 responds to MSP1 and MSP2 (but not MSP3). These results show that crosstalk can occur between MutRS1-harboring *S. mutans* strains and MutRS2-harboring *S. mutans* strains, but the crosstalk is not symmetric. Because no other MutR-class homolog appears in *S. mutans* T8, we believe the crosstalk results from promiscuous MSP1 binding to the MutR2 protein. MSP1 has less steric bulk than MSP2 due to a leucine→phenylalanine substitution at position six (Figure 6B); and this may explain why MSP1 appears to activate MutR2, but MSP2 is unable to activate MutR1.

In *S. mutans* LAR01, MSP1 activates both P_*mutS1*_ and P_*mutS3*_, while MSP3 activates only P_*mutS3*_. Curiously, transcriptional activity of P_*mutS3*_ was unidirectional under MSP1 and MSP3 induction, indicating that these pheromones are not sufficient to activate expression of *CO204_09745* alone. In LAR01, MSP2 failed to activate transcription at any *mutS* promoter. We constructed Δ*mutR* strains for both systems and induced them with either MSP1 or MSP3 (Figure 6D). Activation of P_*mutS1*_ was *mutR1*- and MSP1-dependent. Meanwhile, P_*mutS3*_ activation depends on *mutR1* when induced by MSP1 and depends on *mutR3* when induced by MSP3. MSP1 does not activate P_*mutS3*_ in a Δ*mutR1* strain, nor does MSP3 activate P_*mutS1*_ in the absence of *mutS3*. Thus, the observed intrastrain interaction between MutRS1 and MutRS3 is consistent with a peptide-specific but DNA-promiscuous mechanism.

## 3 Discussion

Discovering the MutRS QS system provides new opportunities to study *S. mutans* biology at the community and species level. It is not surprising that QS controls production of mutacins I, II, III, (and quite probably 1140) given the widespread association of QS systems with bacteriocin operons. This study advances our understanding of lantibiotic regulation in *S. mutans* by confirming that QS control of lantibiotic production extends beyond ComDE regulation of Smb. Our findings suggest that QS may control other lantibiotic mutacins like K8 whose regulatory mechanisms remain uncharacterized. The full extent of the MutRS regulon remains unknown as we limited our experimental scope to genes proximal to the MutRS systems. Our search for MutRS-controlled genes was driven by the MutR box found within the promoter region of *mutS* genes; however, MSP1 activated transcription of P_*mutF*_ in a MutR-dependent manner despite there being no identifiable MutR box (Supplementary Figure S2). These findings suggest that MutR can bind sequences different from the P_*mutS*_ and P_*mutA*_ MutR boxes, or that MutR1 control of immunity gene expression occurs through an indirect mechanism. Either possibility suggests that MutRS systems may interact with genes beyond their immediate genomic context. We were able to activate the MutRS3 system in *S. mutans* LAR01 but were unable to induce expression of the fatty acid biosynthesis operon adjacent to *mutS3*. The link between MutRS3 and its regulon could have been disrupted by mobile genetic elements that surround *mutS3*, making it unclear what MutRS3 controls (if anything). Both the MutRS3 and the MutRS4 systems found in >25% of *S. mutans* strains co-localize with a predicted 3-oxoacyl-ACP synthase gene (Figure 1B, Supplementary Data 1). MutRS3’s regulation of the *CO204_09745*-containing operon may be subject to coregulation requiring an additional set of unknown criteria. The crosstalk between MutRS3 and MutRS1 may indicate that some or all of the MutRS3 functions have been subsumed by MutRS1. Future transcriptomic experiments are required to define MutRS system regulons and elucidate the full extent of MutRS control over physiological processes in *S. mutans*.

Crosstalk between MutRS systems (and Rgg-Shp [46] and ComRS [36], [47]) suggests that interactions between QS systems allow multiple environmental sensors to feed into a common effector network. Within a particular species, crosswalk between homologous QS systems allows for coordination of multiple pathways toward a common goal. Interestingly, our QS database reveals that different species colonizing the same ecological niche contain homologous QS systems. For example, MutRS_*α*_ systems are found in strains of *S. mutans* and its fellow dental pathogen *S. sobrinus*. Thus, QS crosstalk represents a potential mechanism for complementary or antagonistic interspecies interactions. We believe our work argues for a systems-level approach in characterizing the ecological importance of QS in the oral microbiome.

QS systems have historically been grouped by the sequence of their regulator proteins, the orientation of their regulator and pheromone genes, or the sequence of the peptide pheromones. Our database highlights the difficulties with clustering QS systems based on any of these metrics alone. For example, many proteins once classified as Type III Rgg regulators fall within the MutR_*∈*_ subclass. The genomic orientation of the pheromone and regulator genes (Figure 1a) is also an imperfect classifier. While not discussed here, other peptide-based QS systems like RopB-LCP [48] and Tpr-Phr [49] are distinct from Rgg-Shp systems but share the same peptide-regulator orientation. Furthermore, the genomic rearrangement seen with the MutRS3 system in *S. mutans* LAR01 is only one example of several QS systems that are located directly upstream or downstream of mobile genetic element genes. While peptide sequence is a useful tool for identifying QS systems from the same subclass, its utility breaks down when trying to extrapolate across different subclasses within the same regulator family. For decades,the ComRS system in *S. sobrinus* went unidentified because its pheromone lacked the aromatic amino acid motifs found in all previously characterized ComRS systems. (It was finally identified by its cannonical ComRS promoter architecture and a downstream, SDS-aligned open reading frame [36].) For all these reasons, we suggest a multi-criteria approach that combines regulator homology, genomic orientation, peptide sequence, and promoter architecture, as presented here with the MutRS, ComRS, and Rgg-Shp systems. We believe this classification strategy will accelerate the identification of new QS systems and allow us to predict “similar” QS systems that may crosstalk. An accurate classification of QS systems enables deeper exploration of the diversity of biological processes regulated by QS in the oral streptococci.

The number of uncharacterized QS systems in our database suggests that most streptococcal QS systems are unstudied. Even model streptococci like *S. mutans* harbor QS systems that control uncharacterized pathways. In addition to the MutRS systems, our QS database includes many examples of bacteriocin-associated regulators that have not been identified as part of a QS system. For example, the database contains two distinct ComRS-class regulators that appear upstream of bacteriocin operons in many pneumococcal genomes (e.g. SPH_0142 and SPH_2131 in *S. pneumoniae* Hungary19A-6). Both regulators were previously identified in bioinformatic surveys as a generic “XRE-family transcriptional regulator” [50] and a “plcR” homolog [51], respectively. These studies focused on the associated bacteriocins and not the regulators themselves, so they did identify the regulators as part of a ComRS-class QS system, nor did they identify the cognate *comS* genes. In our QS database, we provide a prediction for both the ComS peptide sequence and the P_*comS*_ ComR DNA biding site. Our QS database also contains examples of uncharacterized QS systems associated with non-bacteriocin operons. As mentioned previously, we identified an uncharacterized third ComRS system which is widespread among *S. mutans* strains (Figure 1C). This system, which we call ComRS3, appears to regulate a hybrid polyketide synthase, non-ribosomal peptide synthase (PKS-NRPS) system. While the physiological functions of PKS-NRPS systems vary, a hybrid PKS-NRPS system encoded by the *S. mutans mub* operon produces secondary metabolites that both enhance *S. mutans* peroxide tolerance [52] and participate in inter-kingdom communication with *Candida albincans*, a known symbiotic partner of *S. mutans* associated with poor dental health outcomes [53]. Several elements of the *mub* operon are upregulated in response to XIP1 signaling [54], establishing a precedent for ComRS-regulation of hybrid PKS-NRPS systems in *S. mutans*. The three *S. mutans* MutRS_*α*_ systems, the two *S. pneumoniae* ComRS-class systems, and the *S. mutans* ComRS3 system demonstrate the wealth of unknown QS systems in streptococci. It is unknown how many other QS systems remain undiscovered and what aspects of bacterial physiology they may regulate. We encourage others to use of our database as a roadmap for characterizing new streptococcal QS systems.

## 4 Methods

### 4.1 Bioinformatic survey of streptococcal QS systems

Putative peptide pheromone-based QS regulators were identified using homology-guided BLAST tblastn or blastp searches [55] queried with previously characterized Rgg and ComR regulators. Homology hits for each query were recorded and the genome for the strain containing the homologous regulator was downloaded for manual curation. To ensure a greater diversity in putative QS regulators, weak homology hits (∼30% identity) were re-used as the BLAST query sequence. This process was manually repeated in an iterative manner until the frequency of new regulator sequences was greatly diminished.

Putative pheromone genes were identified using a manually implemented, multiple-criteria approach. All open reading frames (ORFs) longer than six amino acids within the upstream and downstream intergenic regions of an identified regulator were filtered by proximity to nucleotide sequences bearing a close resemblance (Hamming distance less than 2) to the canonical Shine Delgarno Sequence (SDS), AGGAGG. In general, all candidate ORFs within 15 nucleotides of a probable SDS were retained. Next, remaining candidate ORFs were filtered by proximity to nucleotide sequences bearing a close resemblance (Hamming distance less than 2) to the canonical −10 promoter element sequence, TATAAT. In general, all candidate ORFs within 100 nucleotides of a probable −10 promoter element were retained. Due to the fact that many QS regulators self-regulate their own pheromone genes, we hypothesized that a DNA binding motif for each regulator would appear in the promoter region for the correct pheromone gene. The nucleotide regions upstream of −10 elements for the remaining ORFs were grouped according to associated regulator homology and aligned using ClustalOmega [56]. The aligned sequences were then used to identify probable DNA motifs indicative of regulator binding. Using the resulting consensus motif sequence for a given subgroup of highly homologous regulators, we searched the remaining candidate ORF promoter regions for similar nucleotide patterns. If a putative regulator binding motif was identified, its spacing to the ORF’s predicted −10 element was noted. This spacing was hypothesized to be relatively conserved across highly homologous systems, and an “appropriate” regulator binding to −10 element spacing was defined for each group. Candidate ORFs which had a probable SDS, a probable −10 element, and a probable regulator binding site spaced appropriately to said −10 element were predicted as the regulator’s cognate pheromone gene. The identified putative pheromone genes and putative regulator binding sites were then compared across regulator classes. Database entries with no high-confidence pheromone gene prediction were then reassessed using more relaxed search parameters (larger Hamming distances, larger regulator promoter spacing allowances, etc.).

### 4.2 Approximation of QS system prevalence in the *S. mutans* pangenome

A collection of 249 *S. mutans* genomic sequences were downloaded from NCBI and parsed using a custom R script. Next, a fuzzy-matching algorithm allowing for 10% amino acid identity mismatch was used to identify homologs of every distinct *S. mutans* QS regulator identified within our QS database. A regulator was defined as distinct if it exhibited >15% sequence identity dissimilarity to all other regulators within its respective subclass (i.e. all other MutRS_*α*_ regulators). Due to annotation discrepancies with the *comC* and *comD* genes, a strain was said to have the ComCDE system if it contained genes encoding for ComE, ComD, or ComC.

### 4.3 Strains, reagents, and growth conditions

*S. mutans* UA140 and T8 were obtained from Justin Merritt at Oregon Health and Science University; and *S. mutans* LAR01 was provided by Jacqueline Abranches at the University of Florida. Liquid cultures were grown aerobically at 37 °C and 5% CO_2_. Overnight cultures were grown in 50% Todd-Hewitt broth plus 0.5% yeast (THY) and 50% chemically defined medium (CDM) [57] containing 1% glucose. Solid agar plates were made with THY, 1.5% agar, and antibiotic selection when needed: 200 *μ*g/mL kanamycin and/or 4 *μ*g/mL chloramphenicol. Plates were incubated aerobically under 5% CO_2_ at 37 °C. All chemicals were purchased from Sigma Aldrich unless otherwise stated. Enzymes were purchased from New England Biolabs; DNA gene fragments were purchased from Twist Bioscience; and synthetic peptides were purchased from Genscript, Inc. at >90% purity. All strains used in this study are listed in Supplementary Table 1.

### 4.4 DNA manipulation and strain construction

Reporter constructs were synthesized and ligated to a chloramphenicol resistance cassette and homology arms targetting the *mtlD*-*mtlA* locus using Golden Gate assembly [58]. Gene deletion cassettes were similarly obtained by ligating synthesized ∼500-bp homology arm fragments identical to the sequence flanking the target gene to a kanamycin resistance cassette with an orientation opposite the target gene. In cases where synthesis was not possible, DNA components were PCR amplified using NEB Q5 High-Fidelity polymerase and purified using AMPure XP beads (Beckman-Coulter) before use in assembly reactions. The resulting linear DNA fragments were capable of chromosomal recombination and were used to transform *S. mutans*. All synthesized DNA constructs and primers used in this study are listed in Supplementary Table 2.

For *S. mutans* strains UA140 and T8, transformations were performed as follows. Overnight cultures were diluted 1:100 into fresh THY and grown to an OD_600_ of 0.10–0.15. Upon reaching this point, Competence Stimulating Peptide (CSP-18, SGSLSTFFRLFNRSFTQA) was added to a final concentration of 1 *μ*g/mL. The cultures were then incubated for 15 minutes under 5% CO_2_ at 37 °C. Aliquots (500 uL) of culture were then mixed with 5 *μ*l of Golden Gate reaction (∼25–50 ng of DNA) in a 1.5 mL microfuge tube (ThermoFisher Scientific) and placed back in the incubator for 2 hours. Following this incubation step, 200 *μ*l of cells were spread onto THY agar plates supplemented with the appropriate antibiotic. Resulting transformants were screened with colony PCR and verified with Sanger sequencing (Azenta Life Sciences).

*S. mutans* LAR01 transformations were performed as follows. Overnight cultures were pelleted at 3,250×G for 5 minutes. The supernatant was decanted and the cell pellet was resuspended and washed with fresh CDM. The culture was then diluted 1:100 into fresh CDM and grown to a OD_600_ of 0.20. The *comX*-Inducing Peptide 1 (XIP1) was added to a final concentration of 2 *μ*M. Aliquots (500 *μ*l) were then immediately mixed with 5 *μ*l of Golden Gate reaction (∼25–50 ng of DNA) in a 1.5 mL microfuge tube (ThermoFisher Scientific) and placed back in the incubator for 2 hours. All subsequent steps were identical to those performed during *S. mutans* UA140 and T8 transformations.

### 4.5 Fluorescence reporter assay

Overnight cultures of reporter strains and wild-type controls were grown aerobically in 50% THY, 50% CDM at 37 °C with 5 % CO_2_. The next day, the cultures were pelleted at 3,250×G and washed with fresh CDM. All cultures were then diluted 1:50 into 5 mL culture tubes. Once the cultures had reach the appropriate cell density (typically OD_600_ between 0.30 and 0.50), they were split into 2.5 mL control and experimental cultures. The experimental cultures were dosed with MSP (typically to a final concentration of 2 *μ*M) while the control group received an equivalent volume of DMSO. Cultures were mixed by vortexing for 10 seconds and aliquoted (250 *μ*l/well) into a black-walled, 96-well microplate. Each plate also contained eight wild-type controls for background subtraction. Four of these were dosed with MSP and four were dosed with DMSO. The plate was then placed into a Cytation 5 multi-mode plate reader with 5% CO_2_ at 37 °C. Optical density and dual-channel fluorescence were measured every 15 minutes following 10 seconds of shaking for 24 hours. mScarlet-I fluorescence was measured using an excitation wavelength of 564±5 nm and an emission wavelength of 602±5 nm. mNeonGreen fluorescence was measured using an excitation wavelength of 505±5 nm and an emission wavelength of 536±5 nm. At least four biological replicates were performed for each condition.

Fluorescence data was processed as follows. To eliminate the optical density-dependent effects of background fluorescence, non-fluorescent wild-type wells were used to train a LOESS local polynomial regression model as a function of OD_600_. The predictions of this model were then used to subtract background fluorescence from reporter strain wells. The raw, background subtracted fluorescence values were then normalized to the culture’s optical density and passed through a third order Savitzky-Golay filter to dampen noise. Maximum activation was determined using the resulting data.

### 4.6 Deferred antagonism assay

Overnight cultures of each producer strain in 50% THY and 50% CDM were pelleted at 3250×G for 5 minutes. The supernatants were removed and each pellet was resuspended in fresh CDM. An aliquot (80 *μ*l) of the washed overnight culture was then added to 5 mL of fresh THY and incubated until the culture reached mid-log phase. While the strains were growing, fresh induction and control plates were made by spreading 200 *μ*l of 1 mM MSP or 200 *μ*l of DMSO over the surface of THY agar plates, respectively. Upon reaching mid-log phase (OD_600_ between 0.3 and 0.5), 20 *μ*l of each producer strain was spotted onto THY/MSP and THY/DMSO plates (*n*=3–4) and allowed to dry for 20 minutes prior to overnight incubation, agar side up, at 37 °C, 5 % CO_2_. After 18 hours, the plates were exposed to UV light for 20 minutes to ensure subsequent antagonism was not due to actively growing producer strains. Target strain overnight culture (1.25 mL) was added to a 50 mL conical containing 25 mL of 45 °C, soft (0.75%) THY agar. The tube was inverted five times to ensure proper mixing and 5 mL of the suspended target strain was added as an overlay to producer strain plates. After 20 minutes, the plates were transferred agar side down to a 37 °C, 5% CO_2_ incubator. After 24 hours, the diameter of the resulting inhibition rings were measured using digital calipers. The zone of inhibition was defined as the difference in the areas of the inhibition ring and the producer spot.

## Acknowledgements

We thank Suzy Dawid and all members of the Jensen Lab for insightful discussions. This work is supported by the National Institutes of Health under grant GM138210.

