## Supplementary Material for "The MutRS quorum sensing system controls lantibiotic mutacin production in the human pathogen *Streptococcus mutans*"

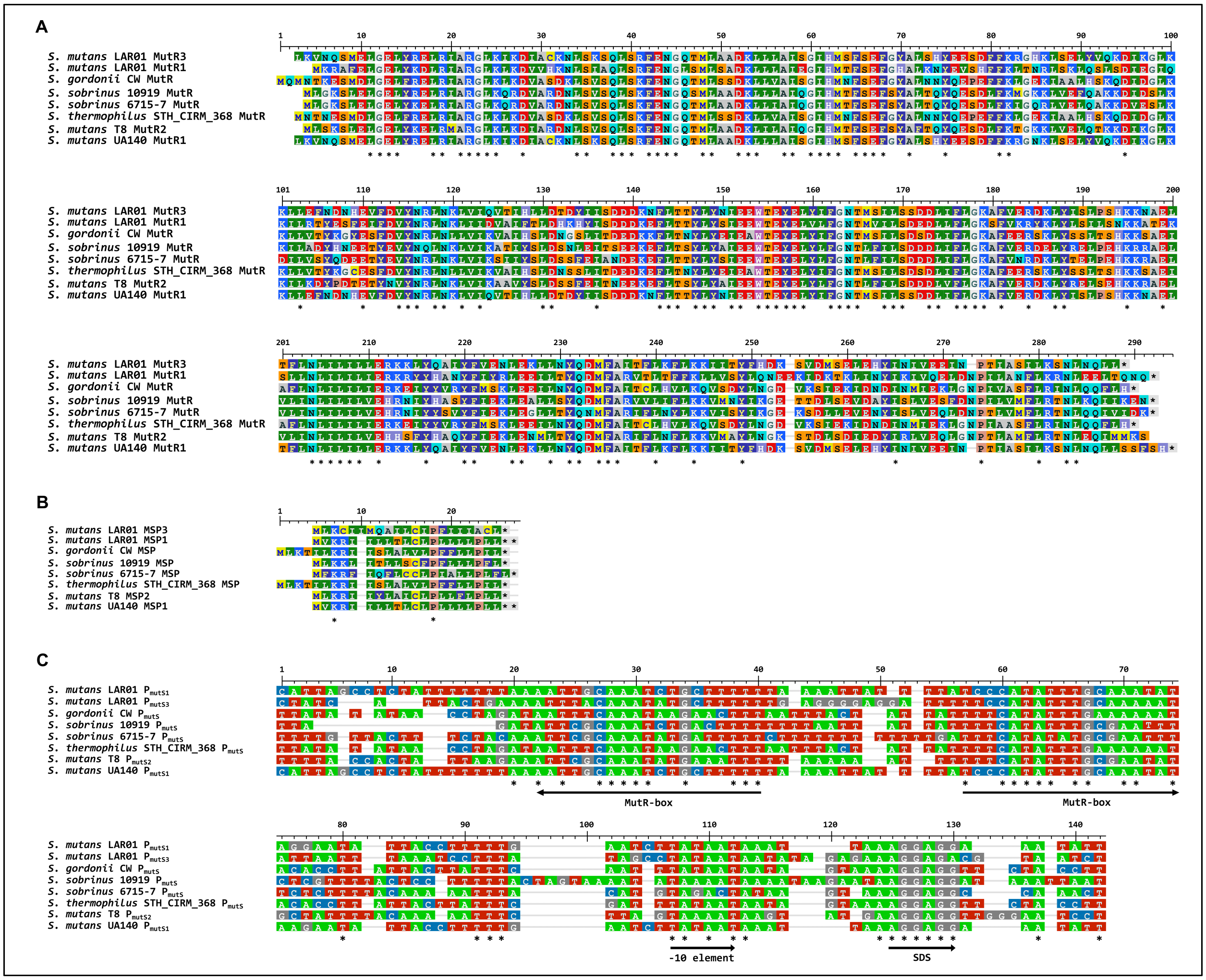


**Figure S1:** Characteristics of representative MutRS$_\alpha$ QS systems. **A.** Clustal Omega alignment of eight representative MutR$_\alpha$ regulators from four different species of streptococci. **B.** Clustal Omega alignment of eight representative MutS$_\alpha$ peptides from four species of streptococci. **C.** Clustal Omega alignment of eight representative P$_{mutS_\alpha}$ nucleotide sequences from four different species of streptococci. * indicate nucleotide identity across all promoter sequences. The left-pointing MutR box is part of the paired promoter (not shown). The paired promoter is P$_{mutA}$ for the *S. mutans* MutRS1 and MutRS2 examples but putatively controls a variety of different genes in the other species shown. The right pointing arrow MutR box belongs to the promoter for *mutS*$_\alpha$, as do the indicated $-10$ element and SDS. Asterisks (*) indicate conserved amino acid identity (**A**–**B**) or nucleotide identity (**C**).

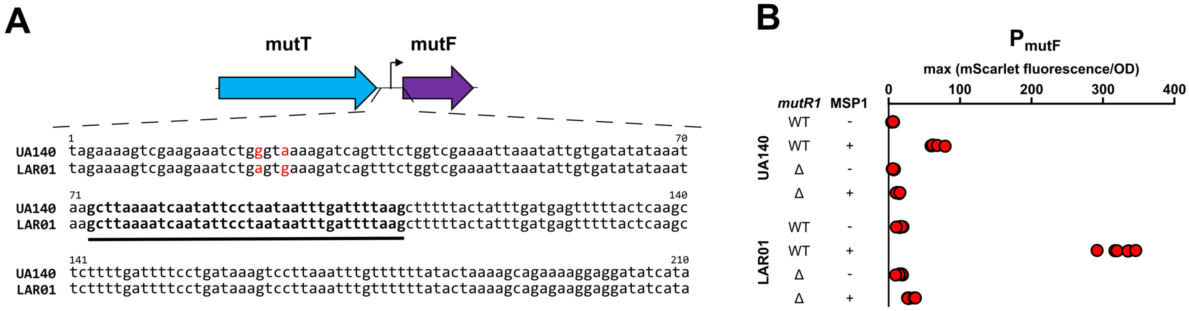


**Figure S2:** MutRS1 systems regulate the expression of lantibiotic immunity genes in *S. mutans* UA140 and LAR01. **A.** Nucleotide sequence of the *mutT*- *mutF* intergenic region. Positions where the nucleotide sequence differs between UA140 and LAR01 are highlighted in red. The promoter architecture in this region is not clear although an imperfect palindromic sequence (bolded and underlined) may indicate the presence of a transcription factor binding site. **B.** Fluorescent reporter experiments demonstrate MSP1-induced and MutR1-dependent activation of $P_{mutF}$ in *S. mutans* UA140 and LAR01. The *mutT–mutF* intergenic region was cloned upstream of the rapidly-maturing *mScarlet-I* fluorescent reporter gene and integrated into the *S. mutans* chromosome at a transcriptionally insulated locus in wild-type and*mutR1* knockout backgrounds. The results indicate the presence of a promoter in this region whose activation depends on MSP1 signaling and the presence of MutR1. However, no MutR box is recognizable in the promoter region, suggesting that the MutR box is more sequence variable than currently understood or that the apparent MutR regulation occurs through an indirect mechanism.

**Supplementary Table 1:** List of strains used in this study

| ID | Strain | Genotype | Description | Source |
|---|---|---|---|---|
| NA | *Streptococcus mutans* UA140 | Wild-type | Parent strain, producer strain for deferred antagonism assays | Justin Merrit, Oregon Health and Science University |
| NA | *Streptococcus mutans* T8 | Wild-type | Parent strain, producer strain for deferred antagonism assays | Justin Merrit, Oregon Health and Science University |
| NA | *Streptococcus mutans* LAR01 | Wild-type | Parent strain, producer strain for deferred antagonism assays | Jacqueline Abranches, University of Florida |
| NA | *Streptococcus mutans* UA159 | Wild-type | Indicator strain for deferred antagonism assays | ATCC |
| NA | Streptococcus *sobrinus* 6715-7 | Wild-type | Indicator strain for deferred antagonism assays | ATCC |
| NA | *Streptococcus gordonii* CH1 | Wild-type | Indicator strain for deferred antagonism assays | ATCC |
| NA | *Lactococcus lactis* subsp. *lactis* ATCC 11454 | Wild-type | Indicator strain for deferred antagonism assays | ATCC |
| NA | *Listeria innocua* ATCC 51742 | Wild-type | Indicator strain for deferred antagonism assays | ATCC |
| RMW46 | *Streptococcus mutans* UA140 | Δ(*mtlD-mtlA*)::PmutSA1_UA140 | Transcriptional reporter strain for UA140 MutRS1 system | This study |
| RMW50 | *Streptococcus mutans* T8 | Δ(*mtlD-mtlA*)::PmutSA2_T8 | Transcriptional reporter strain for T8 MutRS2 system | This study |
| RMW60 | *Streptococcus mutans* LAR01 | Δ(*mtlD-mtlA*)::PmutSA1_LAR01 | Transcriptional reporter strain for LAR01 MutRS1 system | This study |
| RMW107 | *Streptococcus mutans* UA140 | Δ*mutRS1*::*aph3*_KO | Deletion strain for deferred antagonism assays | This study |
| RMW113 | *Streptococcus mutans* T8 | Δ*mutRS2*::*aph3*_KO | Deletion strain for deferred antagonism assays | This study |
| RMW98 | *Streptococcus mutans* LAR01 | Δ*mutRS1*::*aph3*_KO | Deletion strain for deferred antagonism assays | This study |
| RMW108 | *Streptococcus mutans* UA140 | Δ(*mtlD-mtlA*)::PmutSA1_UA140-SynTerm_*cat*, Δ*mutR1::aph3*_KO | Transcriptional reporter strain for UA140 MutRS1 system without MutR1 | This study |
| RMW114 | *Streptococcus mutans* T8 | Δ(*mtlD-mtlA*)::PmutSA2_T8-SynTerm_cat,Δ*mutR2::aph3*_KO | Transcriptional reporter strain for T8 MutRS2 system without MutR2 | This study |
| RMW99 | *Streptococcus mutans* LAR01 | Δ(*mtlD-mtlA*)::PmutSA1_LAR01-SynTerm_cat,Δ*mutR1::aph3*_KO | Transcriptional reporter strain for LAR01 MutRS1 system without MutR1 | This study |
| RMW109 | *Streptococcus mutans* UA140 | Δ(*mtlD-mtlA*)::PmutSA1_UA140-SynTerm_cat,Δ*mutS1::aph3*_KO | Transcriptional reporter strain for UA140 MutRS1 system without MutS1 | This study |
| RMW115 | *Streptococcus mutans* T8 | Δ(*mtlD-mtlA*)::PmutSA2_T8-SynTerm_cat, Δ*mutS2::aph3*_KO | Transcriptional reporter strain for T8 MutRS2 system without MutS2 | This study |
| RMW100 | *Streptococcus mutans* LAR01 | Δ(*mtlD-mtlA*)::PmutSA1_LAR01-SynTerm_cat,Δ*mutS1::aph3*_KO | Transcriptional reporter strain for LAR01 MutRS1 system without MutS1 | This study |
| RMW110 | *Streptococcus mutans* UA140 | Δ(*mtlD-mtlA*)::PmutSA1_UA140-SynTerm_cat, Δ*mutRS1::aph3*_KO | Transcriptional reporter strain for UA140 MutRS1 system without MutRS1 | This study |
| RMW116 | *Streptococcus mutans* T8 | Δ(*mtlD-mtlA*)::PmutSA2_T8-SynTerm_cat, Δ*mutRS2::aph3*_KO | Transcriptional reporter strain for T8 MutRS2 system without MutRS2 | This study |
| RMW101 | *Streptococcus mutans* LAR01 | Δ(*mtlD-mtlA*)::PmutSA1_LAR01-SynTerm_cat,Δ*mutRS1::aph3*_KO | Transcriptional reporter strain for LAR01 MutRS1 system without MutRS1 | This study |
| RMW61 | *Streptococcus mutans* LAR01 | Δ(*mtlD-mtlA*)::PmutSA3_LAR01-SynTerm_cat | Transcriptional reporter strain for LAR01 MutRS3 system | This study |
| RMW223 | *Streptococcus mutans* LAR01 | Δ(*mtlD-mtlA*)::PmutSA3_LAR01-SynTerm_cat,Δ*mutR3::aph3*_KO | Transcriptional reporter strain for LAR01 MutRS3 system without MutR3 | This study |
| RMW239 | *Streptococcus mutans* LAR01 | Δ(*mtlD-mtlA*)::PmutSA3_LAR01-SynTerm_cat,Δ*mutR1::aph3*_KO | Transcriptional reporter strain for LAR01 MutRS3 system without MutR1 | This study |
| RMW219 | *Streptococcus mutans* LAR01 | Δ(*mtlD-mtlA*)::PmutSA1_LAR01-SynTerm_cat,Δ*mutR3::aph3*_KO | Transcriptional reporter strain for LAR01 MutRS1 system without MutR3 | This study |
| RMW142 | *Streptococcus mutans* UA140 | Δ(*mtlD-mtlA*)::PmutF_UA140-SynTerm_cat | Transcriptional reporter strain for UA140 *mutFEG* immunity gene expression | This study |
| RMW144 | *Streptococcus mutans* LAR01 | Δ(*mtlD-mtlA*)::PmutF_LAR01-SynTerm_cat | Transcriptional reporter strain for LAR01 *mutFEG* immunity gene expression | This study |
| RMW143 | *Streptococcus mutans* UA140 | Δ(*mtlD-mtlA*)::PmutF_UA140-SynTerm_cat, Δ*mutR1::aph3*_KO | Transcriptional reporter strain for UA140 *mutFEG* immunity gene expression without MutR1 | This study |
| RMW145 | *Streptococcus mutans* LAR01 | Δ(*mtlD-mtlA*)::PmutF_LAR01-SynTerm_cat, Δ*mutR1::aph3*_KO | Transcriptional reporter strain for LAR01 *mutFEG* immunity gene expression, without MutR1 | This study |

**Supplementary Table 2**: List of synthesized DNA constructs and primers used in this study

| Name | Sequence (5’-3’) | Description |
|---|---|---|
| PmutSA1_UA140 | tcaccaccctggatctacaagtatacggtctcgatagtaaatcaaaaaaatgctaagcctcgtttggcttagcattgcaacttattatttgtaaagttcatc<br>catgcccatgacatctgtgaaggctttttgccattctttgaagttcagctctgtcttactgtgcttcagttctgttttgcggaaaacgtacatgggctgattcttt<br>aagtaattcgcagccattggctttgcaaaggtataagtggtacgggccgtggaacggtagcgtttaccgttaccagtggtgtatgaccacttaaaagt<br>cgaaatgattgttttgtcgttggggtaagtctttttggaacgacaccaatccgcagccgtcagggaatttgtcataacaggtccatctgccgggaatccc<br>gttcccttcacttgggcctcccccttgatatgcgagccctcataagtataacggtaatttactgtaagcgaagctccgtcttcaaattgcatagtacgatg<br>gacctgatatcccgacccgtcaaccatagccgcttgaaagggggacatgccatcggggtacggaaggtactggtggaaaccgtagccgatgtgc<br>ggcactaagatccaaggtgaaaattgcaagtctcccttggtactcttcaggtttaactcctcgtagccgtcgttcggattcccagtaccctggccgacca<br>tatcaaagtccaccccgttgatggatccgaagatatgcaactcatgagttgcgggtaatgaggccatattatcctcctctcctttcgacaccataatatc<br>ctcctttttcatgtgtgtaactgaacgtaaattgttcagtagaagcatatcagaaaacttttaacggtaatctgataatagttattatataattataattatata<br>cttattaagtagtacatctacagataactaatgctagcagaaatatctaagataacaacaaaatctgttgtaatagtaaatcactattactgagtaaac<br>atatatttacaataatataataccaccattagcctctattttttaaaattgcaaatctgctttttaaaattattttatcccatatttgcaaatataagaatattac<br>ctttttgaatcttataataaattaaaggaggaaatattatggtaagtaaaggcgaagcagtcattaaagaatttatgcgctttaaagtacatatggaggg<br>ctctatgaacggtcatgaatttgagatcgagggtgaaggtgagggcagaccatatgaaggaacccagactgccaagcttaaagttaccaaaggtg<br>gtcccttgccctttagctgggatatccttagtccgcaatttatgtatggttctcgtgcattcatcaagcatccggctgatattcccgactattataaacagtctt<br>ttcctgaaggttttaagtgggaacgtgttatgaattttgaggatggtggtgccgtgactgttactcaggatactagcttagaagatggtactttaatctaca<br>aagttaaattgcgtggcaccaactttccaccggatggccccgtgatgcagaaaaagactatgggctgggaggcaagcaccgagagattgtaccc<br>cgaggatggagtgttgaaaggagatatcaaaatggctcttagattaaaagatggtggacgctatttagccgattttaagactacttataaggctaaga<br>aacccgtccaaatgcccggtgcctacaatgtagaccgcaaattggatatcacaagtcataatgaggattatactgtcgtcgaacagtatgagcgca<br>gtgaaggccgccactctactggtggaatggatgagttatataagtaagactggagaccagccgtggagcaaggtctgttgtagt | Duet transcriptional reporter construct containing *S. mutans*-codon optimized *mNeonGreen and mScarlet-I* flanking the UA140 mutA-mutS intergenic region. The *mNeonGreen* fluorescent reporter gene is followed by a strong instrinsic terminator,Term_M102_gp38, derived from *S. mutans* bacteriophage M102 |
| PmutSA2_T8 | tcaccaccctggatctacaagtatacggtctcgatagtaaatcaaaaaaatgctaagcctcgtttggcttagcattgcaacttattatttgtaaagttcatc<br>catgcccatgacatctgtgaaggctttttgccattctttgaagttcagctctgtcttactgtgcttcagttctgttttgcggaaaacgtacatgggctgattcttt<br>aagtaattcgcagccattggctttgcaaaggtataagtggtacgggccgtggaacggtagcgtttaccgttaccagtggtgtatgaccacttaaaagt<br>cgaaatgattgttttgtcgttggggtaagtctttttggaacgacaccaatccgcagccgtcagggaatttgtcataacaggtccatctgccgggaatccc<br>gttcccttcacttgggcctcccccttgatatgcgagccctcataagtataacggtaatttactgtaagcgaagctccgtcttcaaattgcatagtacgatg<br>gacctgatatcccgacccgtcaaccatagccgcttgaaagggggacatgccatcggggtacggaaggtactggtggaaaccgtagccgatgtgc<br>ggcactaagatccaaggtgaaaattgcaagtctcccttggtactcttcaggtttaactcctcgtagccgtcgttcggattcccagtaccctggccgacca<br>tatcaaagtccaccccgttgatggatccgaagatatgcaactcatgagttgcgggtaatgaggccatattatcctcctctcctttcgacaccataataaa<br>acacctcactttattgcttaaaatgtaaaattttgatcaaatattttacatttacagtttaccatctttttttaccactattaagaaattcgcaaatatgaaattttta<br>aaaaatttttcatatttgcgaatatgctattttacaaaaatttcttagtaaaataagtatgaaggaggttgggaatcctatggtaagtaaaggcgaagca<br>gtcattaaagaatttatgcgctttaaagtacatatggagggctctatgaacggtcatgaatttgagatcgagggtgaaggtgagggcagaccatatga<br>aggaacccagactgccaagcttaaagttaccaaaggtggtcccttgccctttagctgggatatccttagtccgcaatttatgtatggttctcgtgcattcat<br>caagcatccggctgatattcccgactattataaacagtcttttcctgaaggttttaagtgggaacgtgttatgaattttgaggatggtggtgccgtgactgtt<br>actcaggatactagcttagaagatggtactttaatctacaaagttaaattgcgtggcaccaactttccaccggatggccccgtgatgcagaaaaaga<br>ctatgggctgggaggcaagcaccgagagattgtaccccgaggatggagtgttgaaaggagatatcaaaatggctcttagattaaaagatggtgga<br>cgctatttagccgattttaagactacttataaggctaagaaacccgtccaaatgcccggtgcctacaatgtagaccgcaaattggatatcacaagtca<br>taatgaggattatactgtcgtcgaacagtatgagcgcagtgaaggccgccactctactggtggaatggatgagttatataagtaagactggagacca<br>gccgtggagcaaggtctgttgtagt | Duet transcriptional reporter construct containing *S. mutans*-codon optimized *mNeonGreen and mScarlet-I* flanking the T8 mutA-mutS intergenic region. The *mNeonGreen* fluorescent reporter gene is followed by a strong instrinsic terminator,Term_M102_gp38, derived from *S. mutans* bacteriophage M102 |
| PmutSA1_LAR01 | tcaccaccctggatctacaagtatacggtctcgatagtaaatcaaaaaaatgctaagcctcgtttggcttagcattgcaacttattatttgtaaagttcatc<br>catgcccatgacatctgtgaaggctttttgccattctttgaagttcagctctgtcttactgtgcttcagttctgttttgcggaaaacgtacatgggctgattcttt<br>aagtaattcgcagccattggctttgcaaaggtataagtggtacgggccgtggaacggtagcgtttaccgttaccagtggtgtatgaccacttaaaagt<br>cgaaatgattgttttgtcgttggggtaagtctttttggaacgacaccaatccgcagccgtcagggaatttgtcataacaggtccatctgccgggaatccc<br>gttcccttcacttgggcctcccccttgatatgcgagccctcataagtataacggtaatttactgtaagcgaagctccgtcttcaaattgcatagtacgatg<br>gacctgatatcccgacccgtcaaccatagccgcttgaaagggggacatgccatcggggtacggaaggtactggtggaaaccgtagccgatgtgc<br>ggcactaagatccaaggtgaaaattgcaagtctcccttggtactcttcaggtttaactcctcgtagccgtcgttcggattcccagtaccctggccgacca<br>tatcaaagtccaccccgttgatggatccgaagatatgcaactcatgagttgcgggtaatgaggccatattatcctcctctcctttcgacaccataatatc<br>ctcctttttcatgtgtgtaactgaacataaattgttcagtagaagcatatcagaaaacttttaacggtaatctgataatagttattatataattataattatata<br>cttattaagtagtacatctacagataactaatgctagcagaaatatctaagataacaacaaaatctgttgtaatagtaaatcactattactgagtaaac<br>atatatttacaataatataataccaccattagcctctattttttaaaattgcaaatctgctttttaaaattattttatcccatatttgcaaatataggaatattac<br>ctttttgaatcttataataaattaaaggaggaaatattatggtaagtaaaggcgaagcagtcattaaagaatttatgcgctttaaagtacatatggaggg<br>ctctatgaacggtcatgaatttgagatcgagggtgaaggtgagggcagaccatatgaaggaacccagactgccaagcttaaagttaccaaaggtg<br>gtcccttgccctttagctgggatatccttagtccgcaatttatgtatggttctcgtgcattcatcaagcatccggctgatattcccgactattataaacagtctt<br>ttcctgaaggttttaagtgggaacgtgttatgaattttgaggatggtggtgccgtgactgttactcaggatactagcttagaagatggtactttaatctaca<br>aagttaaattgcgtggcaccaactttccaccggatggccccgtgatgcagaaaaagactatgggctgggaggcaagcaccgagagattgtaccc<br>cgaggatggagtgttgaaaggagatatcaaaatggctcttagattaaaagatggtggacgctatttagccgattttaagactacttataaggctaaga<br>aacccgtccaaatgcccggtgcctacaatgtagaccgcaaattggatatcacaagtcataatgaggattatactgtcgtcgaacagtatgagcgca<br>gtgaaggccgccactctactggtggaatggatgagttatataagtaagactggagaccagccgtggagcaaggtctgttgtagt | Duet transcriptional reporter construct containing *S. mutans*-codon optimized *mNeonGreen and mScarlet-I* flanking the LAR01 mutA-mutS intergenic region. The *mNeonGreen* reporter gene is followed by a strong instrinsic terminator,Term_M102_gp38, derived from *S. mutans* bacteriophage M102 |
| PmutSA3_LAR01 | tcaccaccctggatctacaagtatacggtctcgatagtaaatcaaaaaaatgctaagcctcgtttggcttagcattgcaacttattatttgtaaagttcatc<br>catgcccatgacatctgtgaaggctttttgccattctttgaagttcagctctgtcttactgtgcttcagttctgttttgcggaaaacgtacatgggctgattcttt<br>aagtaattcgcagccattggctttgcaaaggtataagtggtacgggccgtggaacggtagcgtttaccgttaccagtggtgtatgaccacttaaaagt<br>cgaaatgattgttttgtcgttggggtaagtctttttggaacgacaccaatccgcagccgtcagggaatttgtcataacaggtccatctgccgggaatccc<br>gttcccttcacttgggcctcccccttgatatgcgagccctcataagtataacggtaatttactgtaagcgaagctccgtcttcaaattgcatagtacgatg<br>gacctgatatcccgacccgtcaaccatagccgcttgaaagggggacatgccatcggggtacggaaggtactggtggaaaccgtagccgatgtgc<br>ggcactaagatccaaggtgaaaattgcaagtctcccttggtactcttcaggtttaactcctcgtagccgtcgttcggattcccagtaccctggccgacca<br>tatcaaagtccaccccgttgatggatccgaagatatgcaactcatgagttgcgggtaatgaggccatattatcctcctctcctttcgacaccatttttctcc<br>tccattccattgtttaggtatcagtaatacaagccaaataatctttccatcaatttttagttaaattttaaccgtatcttactttcctatagtatcattaaattctatc<br>attactgaaaatttacaaatatgctttttgaggggaggatttttccatatttgcaaaaatattaatttaaatcctttatagcctataataatatagagaaagg<br>agacgtaatctatggtaagtaaaggcgaagcagtcattaaagaatttatgcgctttaaagtacatatggagggctctatgaacggtcatgaatttgag<br>atcgagggtgaaggtgagggcagaccatatgaaggaacccagactgccaagcttaaagttaccaaaggtggtcccttgccctttagctgggatatc<br>cttagtccgcaatttatgtatggttctcgtgcattcatcaagcatccggctgatattcccgactattataaacagtcttttcctgaaggttttaagtgggaacg<br>tgttatgaattttgaggatggtggtgccgtgactgttactcaggatactagcttagaagatggtactttaatctacaaagttaaattgcgtggcaccaactt<br>tccaccggatggccccgtgatgcagaaaaagactatgggctgggaggcaagcaccgagagattgtaccccgaggatggagtgttgaaaggag<br>atatcaaaatggctcttagattaaaagatggtggacgctatttagccgattttaagactacttataaggctaagaaacccgtccaaatgcccggtgcct<br>acaatgtagaccgcaaattggatatcacaagtcataatgaggattatactgtcgtcgaacagtatgagcgcagtgaaggccgccactctactggtg<br>gaatggatgagttatataagtaagactggagaccagccgtggagcaaggtctgttgtagt | Duet transcriptional reporter construct containing *S. mutans*-codon optimized *mNeonGreen and mScarlet-I* flanking the LAR01 CO204_09745-mutS intergenic region. The *mNeonGreen* fluorescent reporter gene is followed by a strong instrinsic terminator,Term_M102_gp38, derived from *S. mutans* bacteriophage M102 |

| | | |
|---|---|---|
| PmutF_UA140 | tcaccaccctggatctacaagtatacggtctcgatagtagaaaagtcgaagaaatctgggtaaaagatcagtttctggtcgaaaattaaatattgtgat<br>atataaataagcttaaaatcaatattcctaataatttgattttaagctttttactatttgatgagtttttactcaagctcttttgattttcctgataaagtccttaaatt<br>tgtttttatactaaaagcagaaaaggaggatatcataatggtaagtaaaggcgaagcagtcattaaagaatttatgcgctttaaagtacatatggag<br>ggctctatgaacggtcatgaatttgagatcgagggtgaaggtgagggcagaccatatgaaggaacccagactgccaagcttaaagttaccaaag<br>gtggtcccttgccctttagctgggatatccttagtccgcaatttatgtatggttctcgtgcattcatcaagcatccggctgatattcccgactattataaacag<br>tcttttcctgaaggttttaagtgggaacgtgttatgaattttgaggatggtggtgccgtgactgttactcaggatactagcttagaagatggtactttaatcta<br>caaagttaaattgcgtggcaccaactttccaccggatggccccgtgatgcagaaaaagactatgggctgggaggcaagcaccgagagattgtac<br>cccgaggatggagtgttgaaaggagatatcaaaatggctcttagattaaaagatggtggacgctatttagccgattttaagactacttataaggctaa<br>gaaacccgtccaaatgcccggtgcctacaatgtagaccgcaaattggatatcacaagtcataatgaggattatactgtcgtcgaacagtatgagcg<br>cagtgaaggccgccactctactggtggaatggatgagttatataagtaagactggagaccagccgtggagcaaggtctgttgtagt | Transcriptional reporter construct containing *S. mutans*-codon optimized *mScarlet-I* downtream of the UA140 *mutT-mutF* intergenic region. |
| PmutF_LAR01 | tcaccaccctggatctacaagtatacggtctcgatagtagaaaagtcgaagaaatctgagtgaaagatcagtttctggtcgaaaattaaatattgtgat<br>atataaataagcttaaaatcaatattcctaataatttgattttaagctttttactatttgatgagtttttactcaagctcttttgattttcctgataaagtccttaaatt<br>tgtttttatactaaaagcagagaaggaggatatcataatggtaagtaaaggcgaagcagtcattaaagaatttatgcgctttaaagtacatatggag<br>ggctctatgaacggtcatgaatttgagatcgagggtgaaggtgagggcagaccatatgaaggaacccagactgccaagcttaaagttaccaaag<br>gtggtcccttgccctttagctgggatatccttagtccgcaatttatgtatggttctcgtgcattcatcaagcatccggctgatattcccgactattataaacag<br>tcttttcctgaaggttttaagtgggaacgtgttatgaattttgaggatggtggtgccgtgactgttactcaggatactagcttagaagatggtactttaatcta<br>caaagttaaattgcgtggcaccaactttccaccggatggccccgtgatgcagaaaaagactatgggctgggaggcaagcaccgagagattgtac<br>cccgaggatggagtgttgaaaggagatatcaaaatggctcttagattaaaagatggtggacgctatttagccgattttaagactacttataaggctaa<br>gaaacccgtccaaatgcccggtgcctacaatgtagaccgcaaattggatatcacaagtcataatgaggattatactgtcgtcgaacagtatgagcg<br>cagtgaaggccgccactctactggtggaatggatgagttatataagtaagactggagaccagccgtggagcaaggtctgttgtagt | Transcriptional reporter construct containing *S. mutans*-codon optimized *mScarlet-I* downtream of the LAR01 *mutT-mutF* intergenic region. |
| *aph3*_KO | tagttaagtgaataggttcggtcagtaaataatagaaataaaaaatcagacctaagactgatgacaaaaagagaaaattttgataaaatagtcttag<br>aattaaattaaaaagggatctaggatctctttaagaaggagataaatcatggctaaaatgagaatatcaccggaattgaaaaaactgatcgaaaa<br>ataccgctgcgtaaaagatacggaaggaatgtctcctgctaaggtatataagctggtgggagaaaatgaaaacctatatttaaaaatgacggaca<br>gccggtataaagggaccacctatgatgtggaacgggaaaaggacatgatgctatggctggaaggaaagctgcctgttccaaaggtcctgcacttt<br>gaacggcatgatggctggagcaatctgctcatgagtgaggccgatggcgtcctttgctcggaagagtatgaagatgaacaaagccctgaaaagat<br>tatcgagctgtatgcggagtgcatcaggctctttcactccatcgacatatcggattgtccctatacgaatagcttagacagccgcttagccgaattggatt<br>acttactgaataacgatctggccgatgtggattgcgaaaactgggaagaagatactccatttaaagatccgcgcgagctgtatgatttttaaagacg<br>gaaaagcccgaagaggaacttgtcttttcccacggcgacctgggagacagcaacatctttgtgaaagatggcaaagtaagtggctttattgatcttg<br>ggagaagcggcagggcggacaagtggtatgacattgccttctgcgtccggtcgatcagggaggatatcggggaagaacagtatgtcgagctatttt<br>ttgacttactggggatcaagcctgattgggagaaaataaaatattatattttactggatgaattgttttaggtcgacctgcagcgtacg | Positive selection marker cassette used for genetic knockouts. Contains the kanamycin-resistance conferring *aph3* gene under the control of the synthetic constituitve promoter P23. |
| UA140_S1_HA1 | tcaccaccctggatctacaagatagacagttcaacttgctctactcttataatttttcacaaattcagactttaatttcatagcaccaaagccatcaattta<br>caatcaatattgtgatctccttcaacaaggcgaatatttttaacacgtgttccttgttttaaatctttaggagccccccttaactttaagatccttaataagagtg<br>actgtatcaccatcagctaattgtttgccattcgcatcaataacaacaagtccttcttctttctcaacttctgcaggattccattcataagcacattctggac<br>aaaccaatagaataccatcttcatagacatactctgactgacatttaggacaattaggtaaagacatataaactccttttattttgattaccattaacagtt<br>tacctgagtttttatcagttggcaagtcctatcctatcacaacaaaaagctagctgcttgtactgactctatgagttaagtatactgtctaacttatagagtc<br>agtacaagagctagttcttttaattataagagtttcccggactacaacagaccttgctgataaaaataaaaagtctattaaagaataatagactttttata<br>ttatgtagttaagtgatgtaaaacgacggccagtatagactttttatattatgataggacgaatagggagaccggagcaaggtctgttgtagt | Homology arm used for linear construct recombination into UA140 chromosome at mtlD-mtlA locus. The 3' end contains the *L. lactis hisB* strong bidirectional instrinsic terminator and M13F primer binding site |
| UA140_S1_HA2 | tcaccaccctggatctacaaggtctcgactatcacctaattagtttgataatgaagttccgagtttttgtactcgaaccttcaatgaagattttcgttccatca<br>gaaatcccctccttttttattctaaaactactgtaacatttttcattttctaataaaactcaaattttggcacaaaagttttgccaaaatttgttattctaagcgga<br>agaataggatttgacaaagcctgactaagactgactttttatcactaatatttttctcatgttaaaaacttggatatataactttttaacagtttccttaggta<br>atagagacgaaaaaagcagcagcgttcctcaagtttccttttaaacttcaaactgaatgtctcaactctttcatatatttcctctgatggtcaacgttcgtttt<br>accacagcgaaagtagaatttattcttttatgtgactttacggctgcttttactaagctaacacttatctcatacattaaaatattttctgtgaatacaagatttt<br>aaaacagcccaaaaaaaccaaagaaaaatcaagtagatttttctttggtttaagatttagttggagcaaggtctgttgtagt | Homology arm used for linear construct recombination into UA140 chromosome at *mtlD-mtlA* locus |
| T8_S1_HA1 | tcaccaccctggatctacaagatagacagttcaacttgctctactcttataatttttcacaaattcagactttaatttcatagcaccaaagccatcaattta<br>caatcaatattgtgatctccttcaacaaggcgaatatttttaacacgtgttccttgttttaaatctttaggagccccccttaactttaagatccttaataagagtg<br>actgtatcaccatcagctaattgtttgccattcgcatcaataacaacaagtccttcttctttctcaacttctgcaggattccattcataagcacattctggac<br>aaaccaatagaataccatcttcatagacatactctgactgacatttaggacaattaggtaaagacatataaactccttttattttgattaccattaacagtt<br>tacctgagtttttatcagttggcaagtcctatcctatcacaacaaaaagctagctgcttgtactgattctatgagttaagtatactgtctaacttatggaaaa<br>cgcagaacaagagctagttcttttaattataagagtttcccggactacaacagaccttgctgataaaaataaaaagtctattaaagaataatagactttt<br>tatattatgtagttaagtgatgtaaaacgacggccagtatagactttttatattatgataggacgaatagggagaccggagcaaggtctgttgtagt | Homology arm used for linear construct recombination into T8 chromosome at mtlD-mtlA locus.The 3' end contains the *L. lactis hisB* strong bidirectional instrinsic terminator and M13F primer binding site |
| T8_S1_HA2 | tcaccaccctggatctacaaggtctcgactatcacctaattagtttgataatgaagttccgagtttttgtacccgaaccttcaacgaagattttcgttccatc<br>agaaatcccctccttttttattctaaaactactgtaacatttttcattttctaataaaactcaaattttggcaaaacttttgtgccaaaatttattattctaagcgga<br>agaataggatttgacaaagcctgactaagactgactttttatcactaatatttttctcatgttaaaaacttgaatatataactttttaacagtttccttaggta<br>atagagacgaaaaaaagcagcagcattcctcaagtttccttttaaacttcaaactgaatgtctcaactctttcatatatttcctctgatggttaacattcgttt<br>tatcacagcgaaagtggaatttgttcttttatgtgactttacggttgcttttactaagctaacacttatctcatacattaaaatattttctgtgaatacaagatttt<br>aaaacagccaaaaaaaccaaagaaaaatcaagtagatttttctttggtttaagatttagttggagcaaggtctgttgtagt | Homology arm used for linear construct recombination into T8 chromosome at *mtlD-mtlA* locus |
| LAR01_S1_HA1 | tcaccaccctggatctacaagatagacagttcaacttgctctactcttataatttttcacaaattcagactttaatttcatagcaccaaagccatcaattta<br>caatcaatattgtgatctccttcaacaaggcgaatatttttaacacgtgttccttgttttaaatctttaggagccccccttaactttaagatccttaataagagtg<br>actgtatcaccatcagctaattgtttgccattagcatcaataacaacaagtccttcttctttctcaacttctgcaggattccattcataagcacattctggac<br>aaaccaatagaataccatcttcatagacatactccgactgacatttaggacaattaggtaaagacatataaactccttttattttgattaccattaacagtt<br>tacctgagtttttatcagttggcaagtcctatcctatcacaacaaaaagctagctgcttgtactgattctatgagttaagtatactgtctaacttatagaaaa<br>cgcagaacaagagctagttcttttaattataagagtttcccggactacaacagaccttgctgataaaaataaaaagtctattaaagaataatagactttt<br>tatattatgtagttaagtgatgtaaaacgacggccagtatagactttttatattatgataggacgaatagggagaccggagcaaggtctgttgtagt | Homology arm used for linear construct recombination into LAR01 chromosome at mtlD-mtlA locus. The 3' end contains the *L. lactis hisB* strong bidirectional instrinsic terminator and M13F primer binding site |
| LAR01_S1_HA2 | tcaccaccctggatctacaaggtctcgactatcacctaattagtttgataatgaagttccgagtttttgtacccgaaccttcaacgaagattttcgttccatc<br>agaaatcccctccttttttattctaaaactactgtaacatttttcattttctaataaaactcaaattttggcacaaaagttttgtcaaaatttgttatcctaagcgg<br>aagaataggatttgacaaagcctgactaagactgactttttatcactaatatttttctcatgttaaaagcttgaatatataactttttaacagtttccttaggt<br>aatagagacgaaaaaaagcagcagcattcctcaagtttccttttaaatttcaaactgaatgtctcaactctttcatatatttcctctgatggtcaacattcgt<br>tttatcacagcgaaagtggaatttgttcttttatgtgactttacggttgcttttactaagctaacacttatctcatacattaaaatattttctgtgaatacaagattt<br>taaaacagcccaaaaaaaccaaagaaaaatctacttgatttttctttggtttaagatttagttggagcaaggtctgttgtagt | Homology arm used for linear construct recombination into LAR01 chromosome at *mtlD-mtlA* locus |
| SynTerm_cat | tcaccaccctggatctacaaggtctcggactggtcatagctgtttcctgctagcggtcgagcactagataatctagtgctttttatttttactgagtaggac<br>aaatccgccgggagcggatttgaacgttgcgaagcaacggcccggagggtggcgggcaggacgcccgccataaactgccaggcatcaaatta<br>agcccaggcatcaaataaaacgaaaggctcagtcgaaagactgggcctttcgttttatctgttgtttgtcggtgaacgctctctactagagtcacactg<br>gctcaccttcgggtgggcctttctgcgtttatatagttaagtgaataggttcggtcagtaaataataggattttcgttcgtgaatacatgttataataactata | Transciptionally insulated positive selection marker cassette containing the chloramphenicol-resistance |

| | | |
|---|---|---|
| | actaataacgtaacgtaaaatttgtttgatttgtccaaaatatggtataataggtactaatcaaaatagtgaggtaaccaattcggaggcatatcaaatgaactttaataaaattgatttagacaattggaagagaaaagagatatttaatcattatttgaaccaacaaacgacttttagtataaccacagaaattgatattagtgttttataccgaaacataaaacaagaaggatataaattttaccctgcatttattttcttagtgacaagggtgataaactcaaatacagcttttagaactggttacaatagcgacggagagttaggttattgggataagttagagccactttatacaatttttgatggtgtatctaaaacattctctggtatttggactcctgtaaagaatgacttcaaagagttttatgatttatacctttctgatgtagagaaatataatggttcggggaaattgtttcccaaaacacctatacctgaaaatgctttttctctttctattattccatggacttcatttactgggtttaacttaaatatcaataataatagtaattaccttctacccattattacagcaggaaaattcattaataaaggtaattcaatatatttaccgctatctttacaggtacatcattctgtttgtgatggttatcatgcaggattgtttatgaactctattcaggaattgtcagataggcctaatgactggcttttataagtcgacctgcagcgtacgaaaaattgaaaaaatggtggaaacactttttcaattttttgttttattatttaatatttgggaaatattcattctaattggtaatcagattttagaaaacaataaacccttgcataactttctcgtccatatcggaaactactttttattccctttttttcgaccgagattattttgcgataaaatccgattaagatactaggagaccggagcaaggtctgttgtagt | conferring *cat* gene. The 5' end contains a M13R primer binding site; the strong instrinsic terminator,Term_M102_gp20, derived from S. mutans bacteriophage M102; and the bidirectional rrnB/T7 terminator from pDL278. The 3' end contains the *aad9* terminator from pDL278. |
| BlueTag.for | tcaccaccctggatctacaa | Primer used for synthesized construct amplification |
| GreenTag.rev | actacaacagaccttgctcc | Primer used for synthesized construct amplification |
| M13F_Useq.for | tgtaaaacgacggccagt | Sanger sequencing primer |
| M13R_Useq.rev | caggaaacagctatgacc | Sanger sequencing primer |
| UA140_LAR01_dmutR1_HA1.for | tattcctgtgacaaccattgaaacag | Primer for amplification of *mutR1* 5' homology arm in *S. mutans* UA140 and LAR01 |
| UA140_LAR01_dmutR1_HA1.rev | acggctggtctccctattcctttcattataaaggccctcttctg | Primer for amplification of *mutR1* 5' homology arm in *S. mutans* UA140 and LAR01 |
| T8_dmutR2_HA1.for | ctatcgggcaagcatttttgaaag | Primer for amplification of *mutR2* 5' homology arm in *S. mutans* T8 |
| T8_dmutR2_HA1.rev | acggctggtctccctatttaaactctccttaaataaaatcccattatcatag | Primer for amplification of *mutR2* 5' homology arm in *S. mutans* T8 |
| UA140_LAR01_dmutR1_HA2.for | gtatacggtctcggactaagccatcttgataaattttatatctttcatattc | Primer for amplification of *mutR1* 3' homology arm in *S. mutans* UA140 and LAR01 |
| UA140_LAR01_dmutR1_HA2.rev | aaactacccactttactatgagtatct | Primer for amplification of *mutR1* 3' homology arm in *S. mutans* UA140 and LAR01 |
| T8_dmutR2_HA2.for | gtatacggtctcggacttaacctaaggacgataaaaggcac | Primer for amplification of *mutR2* 3' homology arm in *S. mutans* T8 |
| T8_dmutR2_HA2.rev | atgataagaattcagggaataacgg | Primer for amplification of *mutR2* 3' homology arm in *S. mutans* T8 |
| UA140_LAR01_dmutS1_HA1.for | ttgaaagtaaatcaatcaatggaattaggtga | Primer for amplification of *mutS1* 5' homology arm in S. mutans UA140 and LAR01 |
| UA140_LAR01_dmutS1_HA1.rev | acggctggtctccctatgaaaggatgaactgtcactatgtttcattac | Primer for amplification of *mutS1* 5' homology arm in S. mutans UA140 and LAR01 |
| T8_dmutS2_HA1.for | agagtttagctatgccttcactc | Primer for amplification of *mutS2* 5' homology arm in S. mutans T8 |
| T8_dmutS2_HA1.rev | acggctggtctccctatgcagctatatggtaaaatacgttgct | Primer for amplification of *mutS2* 5' homology arm in S. mutans T8 |
| UA140_LAR01_dmutS1_HA2.for | gtatacggtctcggactaatatttcctcctttaatttattataagattcaaaaagg | Primer for amplification of *mutS1* 3' homology arm in S. mutans UA140 and LAR01 |
| UA140_LAR01_dmutS1_HA2.rev | gcctaatggttttctgtacataaaataatcttg | Primer for amplification of *mutS1* 3' homology arm in S. mutans UA140 and LAR01 |
| T8_dmutS2_HA2.for | gtatacggtctcggactttcccaacctccttcatacttattttactaag | Primer for amplification of *mutS2* 3' homology arm in S. mutans T8 |
| T8_dmutS2_HA2.rev | agtacgaatttcatctaatattattccggtaag | Primer for amplification of *mutS2* 3' homology arm in S. mutans T8 |
| dmutR1_check.for | tgaagcggtaacgccag | Colony PCR/Sanger sequencing primer for validating deletion of *mutR1* |

| | | |
|---|---|---|
| | | in *S. mutans* UA140 and LAR01 |
| dmutR2_check.for | aatacgattgccatcgtagacag | Colony PCR/Sanger sequencing primer for validating deletion of *mutR2* in *S. mutans* T8 |
| dmutS1_check.for | cgaagaattgaaaaagtgatataatagcacag | Colony PCR/Sanger sequencing primer for validating deletion of *mutS1* in *S. mutans* UA140 and LAR01 |
| dmutS1_check.rev | aacgtattattttactaactctttgtttaccact | Colony PCR primer for validating deletion of *mutS1* in *S. mutans* UA140 and LAR01 |
| dmutS2_check.for | gcggccgataaacttatcttg | Colony PCR/Sanger sequencing primer for validating deletion of *mutS2* in *S. mutans* T8 |
| dmutS2_check.rev | atctcgccatttgcttgcc | Colony PCR primer for validating deletion of *mutS2* in *S. mutans* T8 |
| SMU_S1_gint_check.for | gcataaaaccaagcagagattcc | Colony PCR/Sanger sequencing primer for validating construct insertion into *S. mutans* *mtlD-mtlA* locus. |
| aph3_KO_v1.for | gtatacggtctcgatagtagttaagtgaataggttcggtcagtaaataatag | Primer for amplification of aph3_KO. Adds BsaI overhang for sense orientation |
| aph_KO_v1.rev | acggctggtctccagtctaattaggtgannnnntannnknannknncgtacgctgcaggtcgac | Primer for amplification of aph3_KO. Adds BsaI overhang for sense orientation and adds molecular barcode |
| aph3_KO_v2.for | gtatacggtctcgatagtaattaggtgannnnntannnknannknncgtacgctgcaggtcgac | Primer for amplification of aph3_KO. Adds BsaI overhang for antisense orientation and adds molecular barcode |
| aph_KO_v2.rev | acggctggtctccagtctagttaagtgaataggttcggtcagtaaataatag | Primer for amplification of aph3_KO. Adds BsaI overhang for antisense orientation |
